# Pervasively thinner neocortex as a transdiagnostic feature of general psychopathology

**DOI:** 10.1101/788232

**Authors:** Adrienne L. Romer, Maxwell L. Elliott, Annchen R. Knodt, Maria L. Sison, David Ireland, Renate Houts, Sandhya Ramrakha, Richie Poulton, Ross Keenan, Tracy R. Melzer, Terrie E. Moffitt, Avshalom Caspi, Ahmad R. Hariri

## Abstract

**Objective:** Neuroimaging research has revealed that structural brain alterations are common across broad diagnostic families of disorders rather than specific to a single psychiatric disorder. Such overlap in the structural brain correlates of mental disorders mirrors already well-documented phenotypic comorbidity of psychiatric symptoms and diagnoses, which can be indexed by a general psychopathology or *p* factor. We hypothesized that if general psychopathology drives the convergence of structural alterations common across disorders then 1) there should be few associations unique to any one diagnostic family of disorders, and 2) associations with the *p* factor should overlap with those for the broader diagnostic families.

**Methods:** Analyses were conducted on structural MRI and psychopathology data collected from 861 members of the population representative Dunedin Study at age 45.

**Results:** Study members with high scores across three broad diagnostic families of disorders (Externalizing, Internalizing, Thought Disorder) exhibited highly overlapping patterns of reduced global and widely distributed parcel-wise neocortical thickness. Study members with high *p* factor scores exhibited patterns of reduced global and parcel-wise neocortical thickness nearly identical to those associated with the three broad diagnostic families.

**Conclusions:** A pattern of pervasively reduced neocortical thickness appears common across all forms of mental disorders and may represent a transdiagnostic feature of general psychopathology. As has been documented with regard to symptoms and diagnoses, the underlying brain structural correlates of mental disorders may not exhibit specificity, the continued pursuit of which may limit progress toward more effective strategies for etiological understanding, prevention, and intervention.

## Introduction

The search for a structural basis of psychopathology in the brain has historically been dominated by case-control studies wherein comparisons are made between groups of individuals with or without a specific psychiatric diagnosis (1). While such studies have reported a multitude of structural brain differences between cases and controls, they have generally failed to identify specific differences in brain structure that are unique to one diagnosis. On the contrary, the results from these studies generally reveal structural features of the brain that are highly conserved across many disorders (e.g., (2)). For example, a meta-analysis of neuroimaging data from 15,892 individuals across six categorical disorders identified transdiagnostic structural deficits within attentional and cognitive control networks (3). A second meta-analysis of data from 14,027 patients identified transdiagnostic structural deficits in multiple cortical regions across eight categorical disorders (4). Thus, it appears that categorical disorders may not systematically differ in their patterns of associated structural brain alterations.

A similar convergence has been documented in research on psychiatric nosology where accumulating evidence demonstrates that sets of disorders/symptoms predictably co-occur and can be captured within broader families of disorders (5,6). For example, depression and anxiety emerge in the same individuals and comprise the Internalizing family; and antisocial behavior and drug abuse emerge in the same individuals and comprise the Externalizing family. Recent work has shown that disorganized thoughts, delusional beliefs, hallucinations, obsessions and compulsions emerge in the same individuals over time and comprise the Thought Disorder family (7). In addition, a single general psychopathology factor, often called the ‘*p’* factor (7), has been identified that robustly captures the shared variance across these three broader families of common mental disorders (8–10). It is thus possible that the aforementioned convergence of structural brain alterations across categorical disorders reflects the pervasive co-occurrence of psychiatric disorders in individuals, which can be indexed by the *p* factor. Indeed, a recent study reported global grey matter volume reductions not only across categorical disorders but also with higher general psychopathology (11).

In the current study, we hypothesized that if general psychopathology drives the convergence of structural alterations common across disorders then 1) there should be few associations unique to any one of three diagnostic families of disorders and 2) associations with the *p* factor should overlap with those for the three diagnostic families. We tested our hypotheses through variability in two structural features of neocortex - cortical thickness and surface area – derived from high-resolution structural MRI data collected from members of the Dunedin Study, which has followed a population-representative birth cohort for five decades. Our focus on neocortex reflects both the preponderance of prior transdiagnostic neuroimaging findings in cortical regions (3,4) as well as the preferential role of cortical circuits in supporting higher-order integrative, executive processes, dysfunctions of which are hypothesized to be a core feature of general psychopathology (10).

## Methods

### Study Design and Population

Participants are members of the Dunedin Study, a longitudinal investigation of health and behavior in a representative birth cohort. Participants (n=1037; 91% of eligible births; 52% male) were all individuals born between April 1972 and March 1973 in Dunedin, New Zealand (NZ), who were eligible based on residence in the province and who participated in the first assessment at age 3 years (12). The cohort represented the full range of socioeconomic status (SES) in the general population of NZ’s South Island and as adults matched the NZ National Health and Nutrition Survey on key adult health indicators (e.g., body mass index, smoking, GP visits) and the NZ Census of citizens of the same age on educational attainment. The cohort is primarily white (93%), matching South Island demographics (12). Assessments were carried out at birth and ages 3, 5, 7, 9, 11, 13, 15, 18, 21, 26, 32, 38, and most recently (completed April 2019) 45 years. The relevant ethics committees approved each phase of the Study and informed consent was obtained from all participants. Of 1037 Study members in the original cohort, 997 were still alive at age 45 years, and 938 took part in the age-45 assessment. Of these, 875 were scanned (441 [50.4%] male). Scanned Study members did not differ from other living Study members on childhood SES or childhood IQ, nor on the *p* factor (Figure S1).

### Assessment of Psychopathology

We have previously described the structure of psychopathology from age 18 up to age 38 years (7); here we use the models extended to include the age 45 data (see Supplement for details). To examine the structure of psychopathology we used ordinal measures that represented the number of the observed symptoms associated with each disorder assessed repeatedly from age 18 to age 45 (Figure S2). Using Confirmatory Factor Analysis, we tested two standard models (10): (a) a correlated-factors model and (b) a hierarchical or bifactor model. Using a correlated-factors model (Figure S3, Model A) we tested three factors representing Externalizing (with loadings from ADHD, conduct disorder, alcohol, cannabis, tobacco, other drug dependence), Internalizing (with loadings from MDE, GAD, fears/phobias, PTSD, eating disorders), and Thought disorders (with loadings from OCD, mania, schizophrenia). The model fit the data well (Table S1) confirming that three correlated factors (i.e., Internalizing, Externalizing, Thought Disorder) explain well the structure of the disorder symptoms.

A hierarchical or bifactor model (Figure S3, Model B) tested the hypothesis that symptom measures reflect both General Psychopathology and three narrower styles of psychopathology. General Psychopathology (labeled *p*) is represented by a factor that directly influences all of the diagnostic symptom factors. In addition, styles of psychopathology are represented by three factors, each of which influences a smaller subset of the symptom items. For example, alcohol symptoms load jointly on the General Psychopathology factor and on the Externalizing style factor. The specific factors represent the constructs of Externalizing, Internalizing, and Thought Disorder apart from General Psychopathology. After identifying a Heywood case, an estimated variance that was negative for one of the lower-order disorder/symptom factors (specifically mania), we respecified the model accordingly (Figure S3, Model B’). This model fit the data well (Table S1). The *p* factor captured how cohort members differ from each other in the variety and persistence of many different kinds of disorders over the adult life course (Figure S4). Cohort members with higher *p* scores experienced a greater variety of psychiatric disorders from early adolescence to midlife (r=.77).

### MRI Data Acquisition & Processing

Each Study member was scanned using a Siemens Skyra 3T scanner equipped with a 64-channel head/neck coil at the Pacific Radiology imaging center in Dunedin, New Zealand. The following scans were collected during scanning: 1) high resolution T1-weighted images, 2) 3D fluid-attenuated inversion recovery (FLAIR) images, and a gradient echo field map (see Supplement for details).

Structural MRI data were analyzed using the Human Connectome Project (HCP) minimal preprocessing pipeline (see Supplement for details). For each Study member the mean cortical thickness and surface area were extracted from each of the 360 cortical areas in the HCP-MPP1.0 parcellation (13). Outputs of the minimal preprocessing pipeline were visually checked for accurate surface generation by examining each Study member’s myelin map, pial surface, and white matter boundaries. Of the 875 Study members for whom data were available, 4 were excluded due to major incidental findings or previous injuries (e.g., large tumors or extensive damage to the brain/skull), 9 due to missing T2-weighted or field map scans, and 1 due to poor surface mapping, yielding 861 datasets for analyses.

### Statistical Analyses

All analyses were conducted in R version 3.4.1 (14). For expository purposes, we scaled Study members’ scores on the three psychopathology factors (Internalizing, Externalizing, Thought Disorder) from the correlated-factor model, as well as *p*, to M=100, SD=15. We used Ordinary Least Squares (OLS) regression to test associations between each of the three psychopathology factors from the correlated-factor model, as well as *p*, and average cortical thickness and total surface area. Next, we used OLS regression to predict cortical thickness and surface area in each of the 360 parcels from the scheme described above (13). We corrected for multiple comparisons across the 360 tests each for *p* factor scores, Internalizing, Externalizing, and Thought Disorder factor scores using a false discovery rate (FDR) procedure (15). Sex was included as a covariate in all analyses. Total surface area and average cortical thickness were not included as covariates in any of the parcel-based analyses because we were interested in examining specific rather than relative regional associations with the psychopathology factor scores as well as regional contributions to general cortex-wide effects. In response to a request from one reviewer, we conducted exploratory whole-brain analyses of grey matter volume using voxel-based morphometry paralleling those described above for our surface-based measures to facilitate comparison with prior research (see Supplement for details).

All analysis code is available at: https://github.com/HaririLab/Publications/tree/master/Romer2019AJP_pCorticalThinning.

### Network Enrichment Analyses

In light of the pervasive transdiagnostic pattern of reduced neocortical thickness associated with general psychopathology identified in our primary analyses described above, we conducted secondary analyses to test whether the strength of the parcel-wise associations between *p* factor scores and cortical thickness were evenly distributed across the cortex or enriched in heteromodal association cortices including frontoparietal and default mode networks (16,17). Specifically, we tested whether the standardized βs describing the parcel-wise associations between *p* factor scores and cortical thickness corresponded with a gradient that situates heteromodal association cortices at one end of a spectrum and unimodal sensory and somatomotor cortices at the other (18). To test correspondence between the two maps, we first parcellated the connectivity gradient into the 360 HCP-MMP1.0 by taking the mean of each parcel. This parcellated gradient was then correlated with the parcel-wise standardized βs for the association between *p* factor scores and cortical thickness. To determine significance, we compared this value to a null distribution generated by spin permutation testing (19,20), in which each of the gradient and standardized β maps were randomly spherically rotated 1000 times and correlated with the other map. Results were considered significant at p<0.05.

## Results

### Cortical Thickness

Global cortical thickness was normally distributed across all Study members (2.556±0.089 mm). Each of the three psychopathology factors from the correlated-factor model was equally and similarly associated with global cortical thickness (**Figure 1A-C**). Study members with high Internalizing scores had thinner global neocortex (*β*=-0.156; 95% CI=-0.223, -0.089; p<0.0001) as did Study members with high Externalizing scores (*β*=-0.164; 95% CI=-0.232, -0.097; p<0.0001), and those with high Thought Disorder scores (*β*=-0.169; 95% CI=-0.234, -0.103; p<0.0001). Consistent with evidence pointing to similar negative associations between each of the three psychopathology factors and global neocortical thickness, Study members with high scores on the transdiagnostic *p* factor also exhibited reduced mean neocortical thickness (*β*=- 0.159; 95% CI=-0.224, -0.093; p<0.0001; **Figure 1D**).

**Figure 1.**
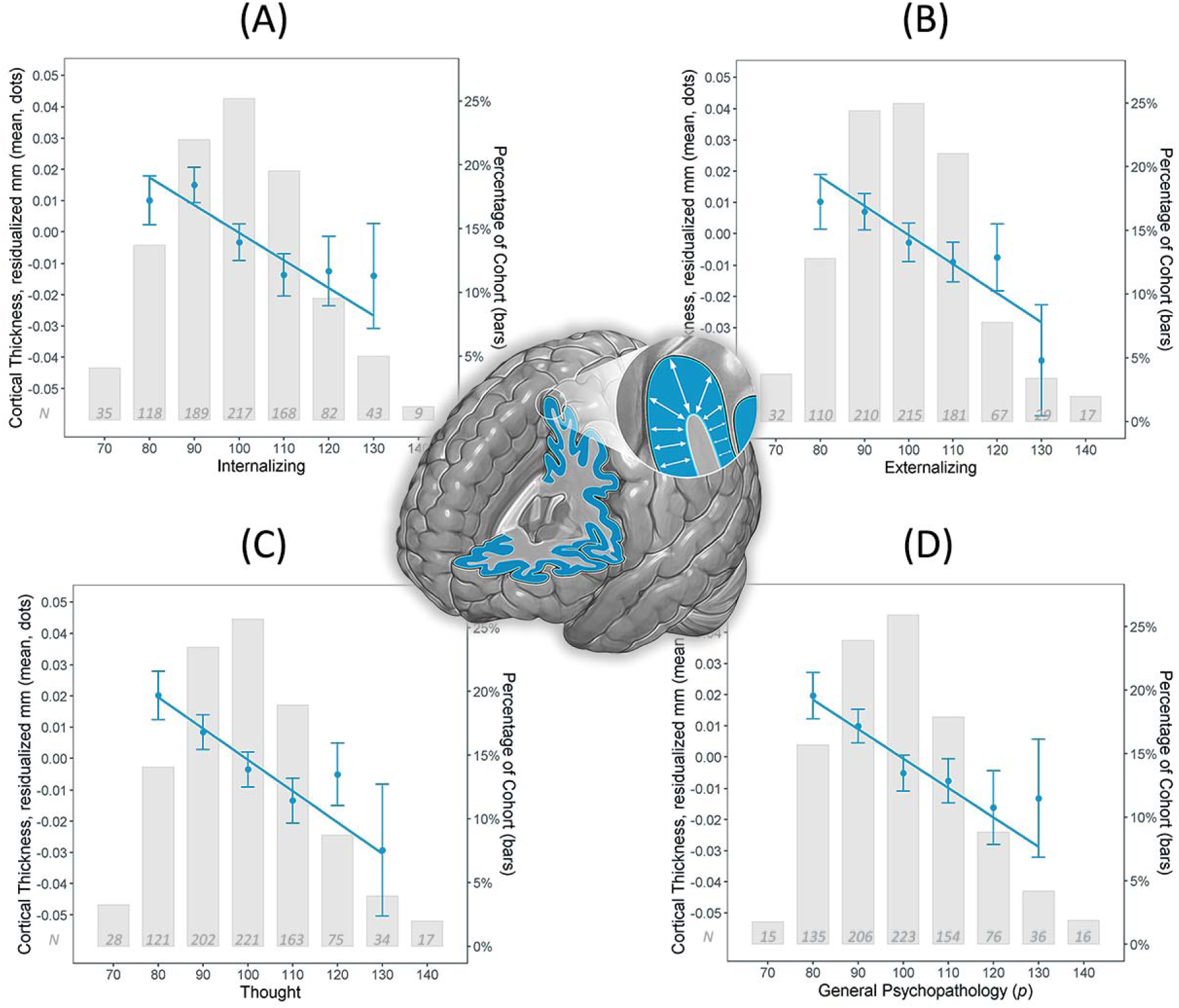
Higher scores across the three broad diagnostic families of disorders (A-C) and general psychopathology as captured by the *p* factor (D) were associated with reduced global cortical thickness. The regression coefficients reported in the text are based on the full distribution of factor scores, which are here clumped for graphing purposes as less than 85, 85 to 95, 95 to 105, 105 to 115, 115 to 125, and greater than 125. The means (dots) and standard errors (bars) show the global cortical thickness in millimeters (residualized on sex) across 861 Study members as a function of diagnostic factor scores. Inset histogram depicts the distribution of factor scores.

In response to requests from reviewers of our original manuscript, we conducted several *post hoc* analyses. First, we examined the potential confounding effects of childhood SES, medical disease, and psychoactive medication use at age 45 years as well as image quality. These analyses revealed that all of the above associations were robust to the inclusion of these covariates (Figure S5). Second, we determined relative effect sizes of global cortical thickness using traditional case-control analyses by comparing four common diagnostic categories observed in the Dunedin Study (i.e., depression, anxiety, substance abuse, schizophrenia) against well-controls (the 147 controls in these analyses represent Study members who have never met diagnostic criteria for any of the mental disorders assessed in the Dunedin Study until age 45; i.e., they have enduring mental health (21). Consistent with our factor-based analyses above, all of these case-control analyses revealed decreased global cortical thickness as a feature of diagnosis (Figure S6). Not surprisingly, the effect was largest for those diagnosed with schizophrenia, which falls at the extreme end of high *p* factor scores. Of note, these effect sizes were larger than the ones observed for the psychopathology factor scores because they involve comparisons between extreme groups (i.e., diagnosed cases versus well-controls that have never met diagnostic criteria for a mental disorder).

Parcel-wise analyses of 360 neocortical areas revealed that high scores on each of the three psychopathology factors were associated with widely distributed patterns of reduced regional cortical thickness (**Figure 2A-C**). High Internalizing scores were associated with significant reductions in cortical thickness in 150 parcels, high Externalizing scores with 171 parcels, and high Thought Disorder scores with 202 parcels (Figure S7 for standardized βs and 95% confidence intervals for all 360 parcels for each factor). Consistent with the global effects above, parcel-wise analyses revealed that high *p* factor scores also were associated with widely distributed reductions in cortical thickness (**Figure 2D**). High *p* factor scores were associated with significant reductions in cortical thickness in 174 parcels (Figure S7). Direct contrasts of the parcel-wise cortical thickness associations with the *p* factor and the Internalizing, Externalizing, and Thought Disorder families of disorders revealed an overlap of 77%, 75%, and 99%, respectively.

**Figure 2.**
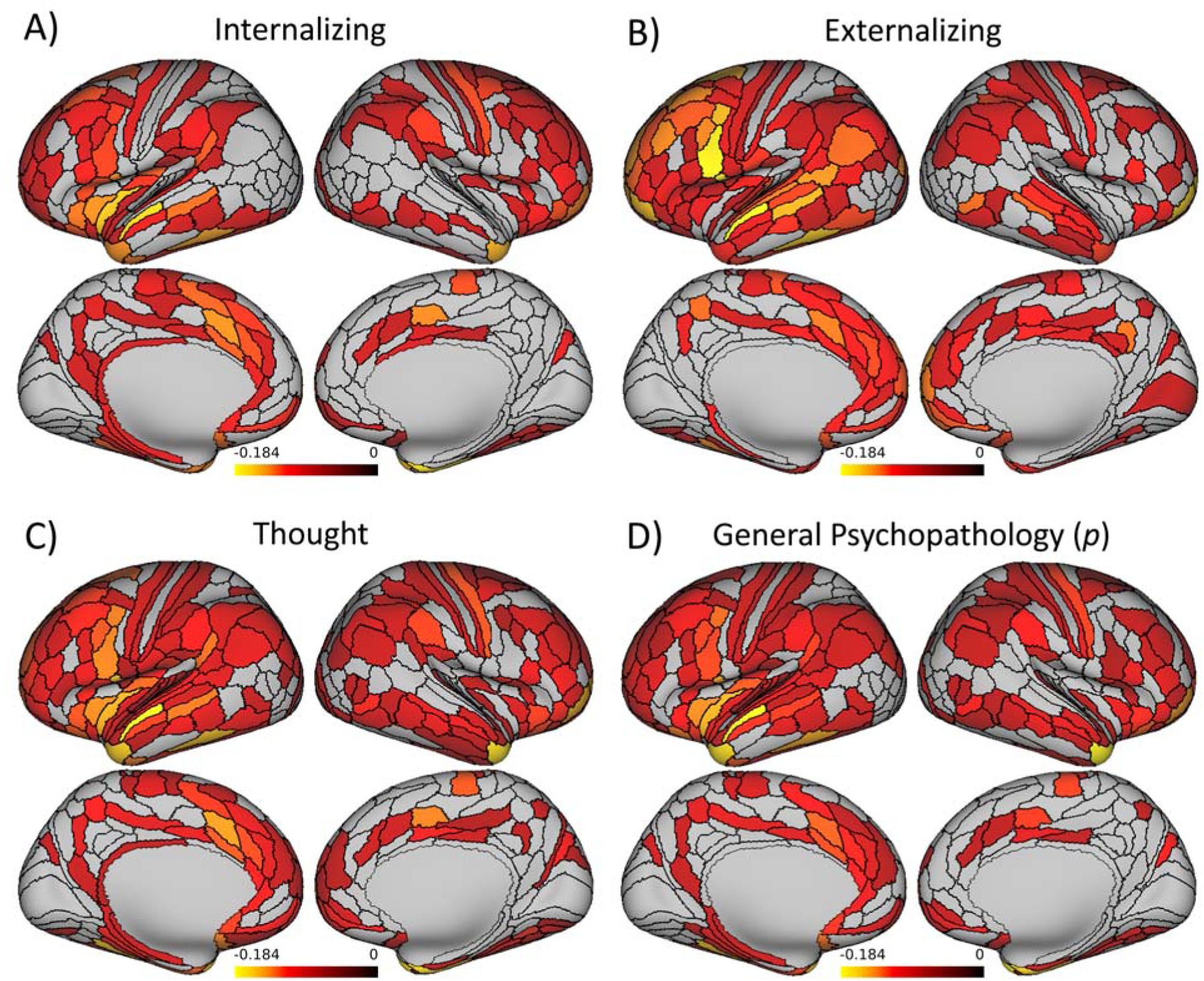
Pervasive and highly overlapping patterns of thinner neocortex were associated with higher scores across the three broad diagnostic families of disorders and general psychopathology as captured by the *p* factor. Statistical parametric maps from parcel-wise analyses are shown to illustrate significant negative associations between cortical thickness and A) Internalizing scores B) Externalizing scores, C) Thought Disorder scores, and D) *p* factor scores. All associations shown are FDR-corrected. Color bars reflect effect sizes (standardized βs).

There were virtually no parcel-wise associations exhibiting reduced cortical thickness that were specific to any of the three broad diagnostic families of disorders (Figure S8). The pervasive and non-specific nature of reduced neocortical thickness was further evident when we compared cortical thickness across all 360 parcels in relation to Internalizing, Externalizing, and Thought Disorder scores, as well as the *p* factor. The high correlations in **Figure 3** (r’s range from .67 to .95) show that those parcels exhibiting reduced cortical thickness among Study members with high scores on one psychopathology factor (e.g., Internalizing) were also reduced among Study members with high scores on the other psychopathology factors (e.g., Externalizing, Thought Disorder). This non-specificity is well captured by evidence that the cortical thickness of the parcels that is reduced among Study members with high scores on any of the three psychopathology factors is also reduced among Study members with high *p* factor scores (r’s range from .72 to .98).

**Figure 3.**
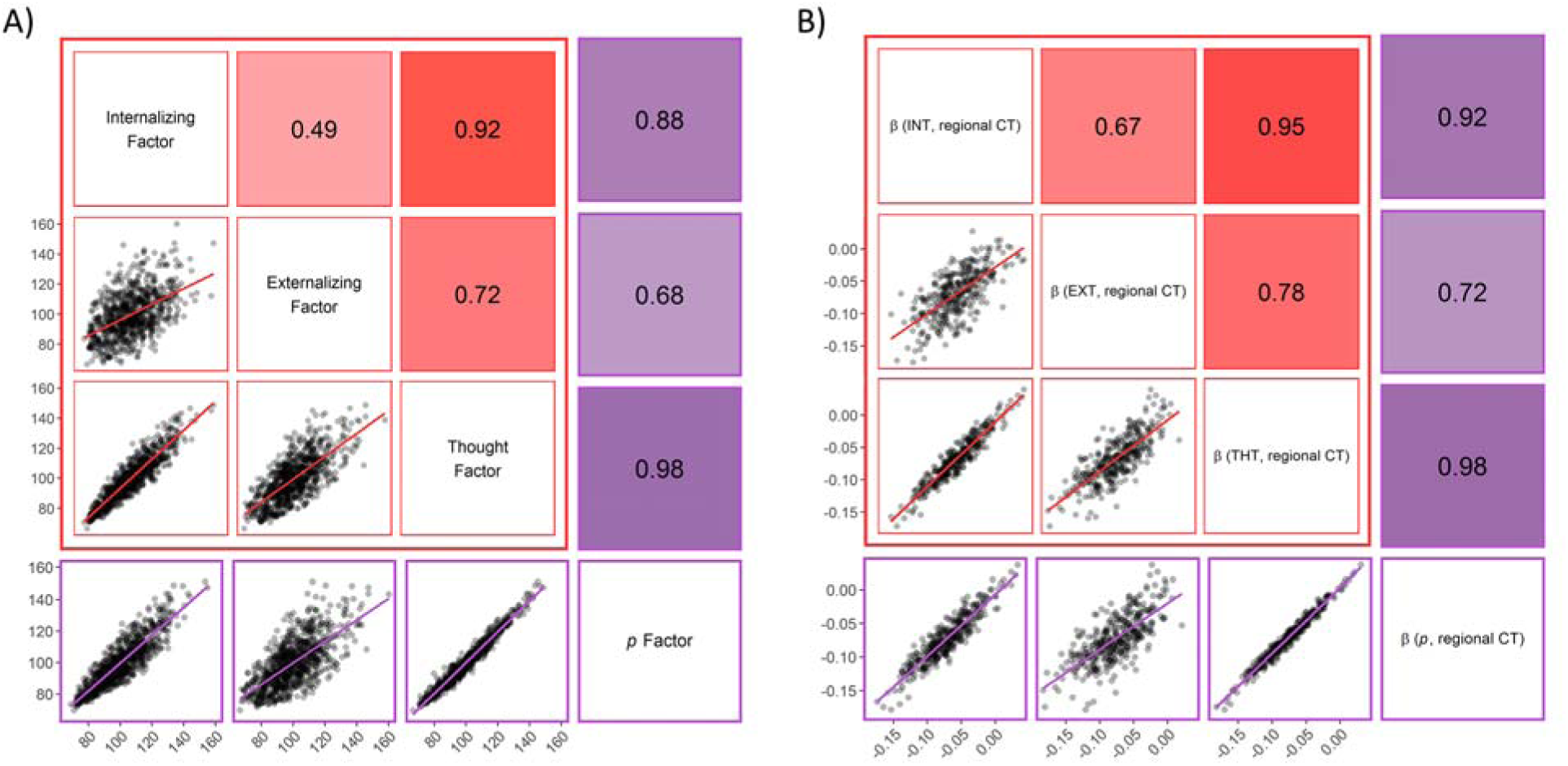
The pervasive pattern of neocortical thinning across all 360 anatomical parcels was highly conserved across the three broader diagnostic families of disorders and the *p* factor, reflecting the relationships between the phenotypes themselves. Correlations between scores for the three factors from the correlated-factors model (red boxes) and between these three factors and the *p* factor from the bifactor model (purple boxes) across all 861 Study members are shown in (A), and between the standardized βs representing the parcel-wise cortical thinning associated with each factor across all 360 parcels are shown in (B). The matrix cells below the diagonal show scatter plots of the associations. The matrix cells above diagonals show their correlations expressed as Pearson’s r. The red and purple regression lines illustrate the slopes of associations between each pair of measures.

### Surface Area

Global surface area was normally distributed across all Study members (185472.1±16347.1 mm^2^). In contrast to the pervasive patterns of reduced cortical thickness found in Study members with higher scores across all factors, a significant reduction in global surface area was only present in Study members scoring higher on the Externalizing factor (β=-0.062, CI=-0.117, -0.007; p=0.026). There were no significant associations between global surface area and scores on either the Internalizing (β=-0.029, CI=-0.084, 0.026; p=0.304) or Thought Disorder factors (β=-0.047, CI=-0.100, 0.007; p=0.087), or the *p* factor (β=-0.044; CI=-0.098, 0.010; p=0.108).

Parcel-wise surface areas were not significantly associated with Internalizing, Externalizing, Thought Disorder, or *p* factor scores. The lack of meaningful parcel-wise associations was further evident when visually examining the standardized βs and 95% confidence intervals for all 360 parcels (Figure S9).

### Network Enrichment

Given that high *p* factor scores were associated with a pervasive transdiagnostic pattern of reduced neocortical thickness, we conducted exploratory analyses to test whether larger than expected associations between *p* factor scores and reduced cortical thickness were present in heteromodal association cortices including the frontoparietal and default mode networks. Spatial permutation testing revealed a significant spatial correspondence between the strength of parcel-wise associations between cortical thickness and *p* factor scores (i.e., standardized β*s*) and a cortical gradient spanning from heteromodal association cortices, on one end, to unimodal sensory and somatomotor cortices, on the other end (**Figure 4**). Specifically, parcels encompassing heteromodal association cortices tended to have larger negative associations between cortical thickness and *p* factor scores than parcels encompassing unimodal sensory and somatomotor cortices (Spearman’s ρ=-0.203, p=0.049).

**Figure 4.**
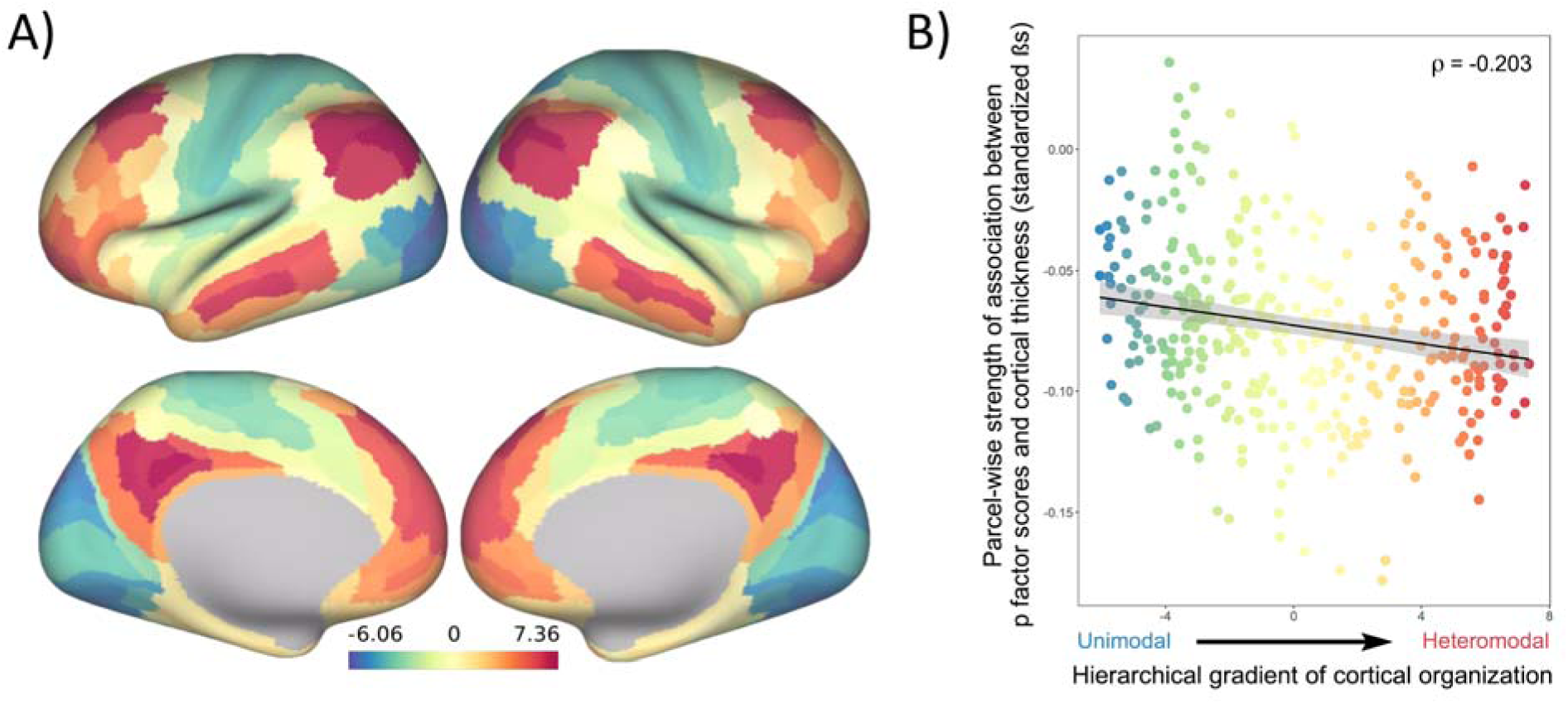
Correspondence of parcel-wise associations between higher *p* factor scores and reduced cortical thickness and a cortical gradient spanning from heteromodal association cortices, including frontoparietal and default mode networks, on one end, to unimodal sensory and somatomotor cortices, on the other end (A; adapted from (18)). Spatial permutation testing revealed that parcels encompassing heteromodal association cortices tended to have larger negative associations (standardized βs) between cortical thickness and *p* factor scores than parcels encompassing unimodal sensory and somatomotor cortices (B; Spearman’s ρ = -0.203, p = 0.049).

## Discussion

The current analyses suggest that there is little specificity in the brain structural correlates of mental disorders. Rather, a non-specific and pervasive pattern of thinner neocortex appears to be a transdiagnostic feature of general psychopathology as indexed by the *p* factor. Consistent with our hypotheses that general psychopathology (*p*) drives the convergence of structural alterations common across disorders, we found 1) very few associations unique to the Internalizing, Externalizing, or Thought Disorder diagnostic families and 2) that associations with the *p* factor highly overlapped with those for the three diagnostic families. The pervasive and transdiagnostic nature of these associations is consistent with studies revealing that most structural brain differences are not unique to categorical mental disorders but rather shared across disorders (3,4,11).

The current findings help to refine the interpretation of recent studies that have identified widely-distributed reductions in neocortical gray matter volumes associated with higher general psychopathology (10,11,22). Specifically, our findings suggest that these prior associations may be driven by reduced neocortical thickness and not surface area, which together comprise gray matter volume. Unlike the broad mapping of reduced global and parcel-wise cortical thickness onto psychopathology, there were no significant associations with parcel-wise neocortical surface area. While there was a significant association with reduced global surface area, this was restricted to higher scores only on the Externalizing factor. This is consistent with prior research showing that Externalizing disorders tend to load least strongly on the *p* factor (7,23). As we did not hypothesize this effect, and it could arise from chance, we refrain from extensive discussion. It raises the question of whether this specific association reflects early life vulnerability to externalizing disorders due to differences in the developing brain (e.g., due to genetics or maltreatment) versus harm done to the brain from an externalizing lifestyle (e.g., due to drug use, sexually transmitted diseases, violent injuries including concussions)(24,25). Regardless, the effect sizes associated with either global or parcel-wise surface area, irrespective of statistical significance, were an order of magnitude smaller than those observed for associations with global and parcel-wise cortical thickness.

Thus, the general pattern of associations suggests that reduced cortical thickness and not surface area is a transdiagnostic feature of mental disorders reflected in general psychopathology. Consistent with this pattern, a recent cross-disorder genome wide association study identified four common variants predicted to impact the function of radial glia and interneurons critical for the development of cortical layers, which are reflected in cortical thickness (26). That said, the cytological and histological basis of cortical thickness remains unclear. MRI-derived estimates of cortical thickness do accurately reflect the width of the cortical mantle as determined postmortem (27). Differences in cortical thickness, however, are unlikely to reflect numbers of neurons or loss of neurons over time (28). Rather, the thickness of the cortex likely reflects a combination of neuron size (i.e., degree of shrinkage) and dendritic arborization (i.e., degree of branching, spine density).

A thinner neocortex has been associated with a host of negative outcomes across the lifespan. For example, a thinner neocortex has been associated with lower intelligence in midlife (29) and is a feature of older brain age relative to chronological age, which is accompanied by greater cognitive impairment (30). A thinner cortex has also been observed in categorical mental disorders spanning the Internalizing, Externalizing, and Thought Disorder diagnostic families (e.g., (31–33)). Meanwhile, accelerated thinning has been associated with worsening daily functioning and other symptoms in Alzheimer’s disease (34,35). Our secondary analyses of network enrichment are relevant here as they highlight preferential associations between higher *p* factor scores and reduced cortical thickness of parcels falling within heteromodal association cortices, including the frontoparietal and default mode networks, supporting higher cognitive processes and executive functions (36). Thus, while a pervasive pattern of reduced neocortical thickness may be a transdiagnostic feature of general psychopathology, it is possible that specific reductions in heteromodal association cortices (i.e., frontoparietal and default mode networks) may feature in the disordered form and content of thought hypothesized to represent the core of the *p* factor (10,37). This is consistent with prior transdiagnostic research indicating alterations within frontoparietal and default mode networks supporting executive control and self-referential processes across diagnostic categories (2,4,36,37). Further evidence comes from parallel neuroimaging studies implicating structural alterations within a cerebello-thalamo-cortical circuit supporting executive functions in the expression of the *p* factor (38–40).

### Limitations

One limitation of the current study is the availability of only a single, cross-sectional assessment of brain structure in midlife. Unfortunately, we have only recently been able to collect neuroimaging data in the Dunedin Study and we are unable to examine temporal order in the data. It remains to be determined if a higher burden of psychopathology contributes to a pervasively thinner neocortex or if a pervasively thinner cortex contributes to a higher burden of psychopathology. Longitudinal collection of neuroimaging data beginning in early childhood and continuing throughout life is needed to address this question. Comparisons with other cross-sectional studies, as well as longitudinal data, are necessary to determine the extent to which the patterns observed here--at age 45 years--generalize across development or the extent to which associations between psychopathology and brain structure change across the lifespan. Second, while the Dunedin Study is a population representative birth cohort free of the selection biases often present in neuroimaging research, it is a predominantly white cohort born in the 1970s in one part of the world; therefore, results will require replication in other samples from other ethnic groups and countries.

### Conclusions

Analyses of data collected from a large population-representative birth cohort in midlife revealed that a pervasively thinner neocortex is a transdiagnostic feature of mental disorders reflecting general psychopathology as indexed by the *p* factor. In addition to furthering the utility of the *p* factor in capturing the shared phenotypic features of common mental disorders, our findings reinforce the value of a transdiagnostic approach, ideally including explicit modeling of general psychopathology, in future neuropsychiatric research. More broadly, our findings underscore that a continued search for specificity amongst mental disorders may not only be elusive but also likely counterproductive in addressing existing gaps in etiology, treatment, and prevention research.

## Acknowledgements

This research was supported by National Institute on Aging grants R01AG032282 and R01AG049789, and UK Medical Research Council grant MR/P005918/1. Additional support was provided by the Jacobs Foundation. A.L.R. and M.L.E. received support from the National Science Foundation Graduate Research Fellowship under Grant No. DGE-1106401 and DGE-1644868, respectively. T.R.M received support from a Sir Charles Hercus Career Development Fellowship from the New Zealand Health Research Council (17/039). The Dunedin Multidisciplinary Health and Development Research Unit was supported by the New Zealand Health Research Council and New Zealand Ministry of Business, Innovation, and Employment (MBIE). We thank members of the Advisory Board for the Dunedin Neuroimaging Study, Dunedin Study members, unit research staff, Pacific Radiology staff, and Study founder Phil Silva, PhD, University of Otago.

The authors declare no competing interests.

