## Supplemental Information for "Pervasively thinner neocortex as a transdiagnostic feature of general psychopathology"

**Transdiagnostic Feature of General Psychopathology**

**Supplemental Materials**

TABLE S2. Summary statistics and anatomical labels for significant clusters in the VBM analyses of grey matter volume correlates of the four factor scores depicted in Figure S10……………………………………………............................... 35

*Participant Attrition Analyses*

We conducted an attrition analysis using childhood intelligence quotient (IQ), childhood socioeconomic status (SES), and *p* factor scores (**Figure S1**).

No significant differences in childhood IQ were found between the full cohort, those still alive, those seen at Phase 45 or those scanned at Phase 45. Those who were deceased by the Phase 45 data collection had significantly lower childhood IQ’s than those who were still alive (t = 2.09, p = 0.04).

No significant differences were found between the full cohort, those deceased, those alive, those seen at Phase 45 or those scanned at Phase 45 on childhood SES.

No significant differences in *p* factor scores were found between the full cohort, those still alive, those seen at Phase 45 or those scanned at Phase 45. Those who were deceased by the Phase 45 data collection had significantly higher *p* factor scores than those who were still alive (t = 2.86, p = 0.004).

*Assessing Psychopathology*

Mental disorders are disturbances in thought, behavior, and emotion that interfere with or limit social, family, educational, or work activities. In the Dunedin Study, these were identified according to the criteria of the Diagnostic and Statistical Manual of Mental Disorders (DSM).

Psychiatric interviews were carried out by health professionals, not lay interviewers. At ages 18, 21, 26, 32, 38, and 45, interviews were carried out with the Diagnostic Interview Schedule (1,2). The following disorders were assessed (**Figure S2**): Externalizing (ADHD, Conduct Disorder, Alcohol Dependence, Tobacco Dependence, Cannabis Dependence, Other Drug Dependence), Internalizing (Generalized Anxiety Disorder, Depression, Fears [including Social Phobia, Simple Phobia, Agoraphobia, Panic Disorder], Eating Disorders [including Bulimia and Anorexia], PTSD), and Thought disorders (Obsessive-Compulsive Disorder, Mania, Schizophrenia). To allow the study of comorbidity, multiple diagnoses could be assigned to a participant at once. However, DSM exclusionary criteria were applied (e.g., hallucinations better explained by drug use were not counted toward schizophrenia; generalized anxiety disorder was not diagnosed if the anxiety stemmed solely from fear about public speaking). The diagnoses were made using computerized algorithms matching the DSM criteria, and additionally requiring self-reported impairment ratings. For disorders where self-reports can be compromised by lack of insight (such as schizophrenia, mania), we also turned to information from additional sources, such as interviews with parents, systematic questionnaires mailed to informants who know the study member well (present data for 97% of the cohort), standardized clinical staff ratings (of observed behavior, such as poor grooming or bizarre speech, during the day of assessment), and medical records for each cohort member from the New Zealand national health system. In the case of schizophrenia and mania, narrative dossiers of symptoms were reviewed by experienced psychiatrists to achieve diagnostic consensus. These details are reported in our previous publications (see, e.g. (3)).

At ages 18 and 21, diagnoses were made according to DSM-III-R (3); at ages 26 , 32, and 38, according to DSM-IV (4); at age 45 according to the now-current DSM-V (5) (with the exception of substance-dependence disorders which were diagnosed according to DSM-IV, given that DSM-V dropped the distinction between abuse and dependence).

*Modelling the Structure of Psychopathology*

We have previously described the structure of psychopathology up to age 38 years (6); here we extend these models to include the age 45 data.

To evaluate the structure of psychopathology we used data from 6 adult assessments, carried out at ages 18, 21, 26, 32, 38, and 45 years. We studied DSM-defined symptoms of the following disorders that were repeatedly assessed in our longitudinal study: ADHD, Conduct Disorder, Alcohol Dependence, Cannabis Dependence, Dependence on Hard Drugs, Tobacco Dependence (assessed with the Fagerström Test for Nicotine Dependence (7)), Depression, Generalized Anxiety Disorder, Fears/Phobias (Social Phobia, Simple Phobia, Agoraphobia, Panic Disorder), PTSD, Eating Disorders (Anorexia, Bulimia), Obsessive-Compulsive Disorder, Mania, and positive and negative Schizophrenia symptoms. Ordinal measures represented the number of the observed DSM-defined symptoms associated with each disorder. Fears/phobias were assessed as the count of diagnoses for simple phobia, social phobia, agoraphobia, and panic disorder that a study member reported at each assessment. Of the 14 disorders, 6 were not assessed at every occasion, but each disorder was measured at least three times (**Figure S2**). Of the original 1,037 study members, we included 1,000 study members who had symptom count assessments for at least one age (845 study members had present symptom counts for all six assessments, 90 for five, 30 for four, 13 for three, and 14 for two). The 37 excluded study members comprised those who died (N=13) or left the Study (N=21) before age 18 or who had such severe developmental disabilities (N=3) that they could not be interviewed with the Diagnostic Interview Schedule.

Using Confirmatory Factor Analysis (CFA), we tested two standard models that are frequently used to examine the structure of psychopathology (8): (a) a correlated-factors model and (b) a hierarchical or bifactor model. In CFA, latent continuous factors are hypothesized to account for the pattern of covariance among observed variables. Our CFAs were run as multitrait-multimethod models. In these models, observed variables represented each of the disorders with a symptom scale at each assessment age (e.g., alcohol dependence was measured with a symptom scale at ages 18, 21, 26, 32, 38, and 45). Each model also included method/state factors designed to pull out age- and assessment-related variance (e.g., interviewer effects, mood effects, and age-specific vulnerabilities) that was uncorrelated with trait propensity toward psychopathology. Because symptom-level data are ordinal and have highly skewed distributions, we used polychoric correlations when testing our models. Polychoric correlations provide estimates of the Pearson correlation by mapping thresholds to underlying normally distributed continuous latent variables that are assumed to give rise to the observed ordinal variables. All CFA analyses were performed in Mplus version 8.3 (9) using the weighted least squares means and variance adjusted (WLSMV) algorithm. The WLSMV estimator is appropriate for categorical and nonmultivariate normal data and provides consistent estimates when data are missing at random with respect to covariates (10). We assessed how well each model fit the data using the chi-square value, the comparative fit index (CFI), the Tucker-Lewis index (TLI), and the root-mean-square error of approximation (RMSEA). CFI values greater than .95 and TLI values greater than 0.95 indicate good fit; RMSEA scores less than .05 are considered good (11).

The correlated-factors model (**Figure S3, Model A**) tests the hypothesis that there are latent trait factors, each of which influences a subset of the diagnostic symptoms. We tested three factors representing Externalizing (with loadings from ADHD, conduct disorder, alcohol, cannabis, tobacco, and other drug dependence), Internalizing (with loadings from MDE, GAD, fears/phobias, PTSD, and eating disorders), and Thought Disorder (with loadings from OCD, mania, and schizophrenia). The model fit the data well: χ^2^(2465, N=1,000) = 4082.230, CFI = .933, TLI = .929, RMSEA = .026, 90% confidence interval (CI) = [.024, .027]. As shown in **Table S1**, loadings on the three specific factors were all positive, generally high (all ps < .001), and averaged .790—Externalizing: average loading = .743; Internalizing: average loading = .814; Thought Disorder: average loading = .844. Correlations between the three factors were all positive and ranged from .420 between Internalizing and Externalizing to .847 between Internalizing and Thought Disorder. Thus, this model confirmed that three correlated factors (i.e., Internalizing, Externalizing, and Thought Disorder) explain well the structure of the disorder symptoms examined across 27 years of adulthood.

The hierarchical or bifactor model (**Figure S3, Model B**) tests the hypothesis that the symptom measures reflect both General Psychopathology and three narrower styles of psychopathology. General Psychopathology (labeled *p* in **Figure S3, Model B**) is represented by a factor that directly influences all of the diagnostic symptom factors. In addition, styles of psychopathology are represented by three factors, each of which influences a smaller subset of the symptom items. For example, alcohol symptoms load jointly on the General Psychopathology factor and on the Externalizing style factor. The specific factors represent the constructs of Externalizing, Internalizing, and Thought Disorder over and above General Psychopathology. Model B had a Heywood case, an estimated variance that was negative for one of the lower-order disorder/symptom factors (specifically, mania), suggesting this was not a valid model. Inspection of the results revealed the source of the convergence problem. Specifically, the Thought Disorder factor was subsumed in *p*; that is, in the hierarchical model, symptoms of OCD, mania, and schizophrenia loaded very highly on *p*, but unlike symptoms of Externalizing and Internalizing, they could not form a separate Thought Disorder factor independently of *p*. We respecified the model accordingly, depicted in **Model B′ Figure S3**. This model fit the data well: χ^2^(2457, N=1,000) = 3695.364, CFI = .949, TLI = .945, RMSEA = .022, 90% CI [.021, .024]. As shown in **Table S1**, loadings on the General factor (*p*) were all positive, generally high (all ps < .001), and averaged .612; the highest standardized loadings were for mania (.976), schizophrenia (.865), PTSD (.860), and OCD (.772). **Figure S4** shows that the *p* factor captures how cohort members differ from each other in the variety and persistence of many different kinds of disorders over the adult life course. Cohort members with higher *p* scores experienced a greater variety of psychiatric disorders from adolescence to midlife (r=.76).

*MRI Acquisition Parameters*

MP-RAGE: TR = 2400 ms; TE = 1.98 ms; 208 sagittal slices; flip angle, 9°; FOV = 224 mm; matrix = 256×256; slice thickness = 0.9 mm with no gap (voxel size = 0.9×0.875×0.875 mm); and total scan time = 6 min and 52 s.

3D fluid-attenuated inversion recovery (FLAIR): TR = 8000 ms; TE = 399 ms; 160 sagittal slices; FOV = 240mm; matrix = 232×256; slice thickness = 1.2 mm (voxel size = 0.9×0.9×1.2 mm); and total scan time = 5 min and 38 s.

Gradient echo field map: TR = 712 ms; TE = 4.92 and 7.38 ms; 72 axial slices; FOV = 200mm; matrix = 100×100; slice thickness = 2.0 mm (voxel size = 2x2x2 mm); and total scan time = 2min and 25 s.

*MRI Data Processing*

T1-weighted and FLAIR images were processed through the PreFreeSurfer, FreeSurfer, and PostFreeSurfer pipelines. T1-weighted and FLAIR images were corrected for readout distortion using the gradient echo field map, coregistered, brain-extracted, and aligned together in the native T1 space using boundary-based registration (12). Images were then processed with a custom FreeSurfer recon-all pipeline that is optimized for structural MRI with higher resolution than 1 mm isotropic. Finally, recon-all output were converted into CIFTI format and registered to common 32k_FS_LR mesh using MSM-sulc (13).

*Voxel-Based Morphometry (VBM)*

Regional gray matter volumes were determined using the unified segmentation and DARTEL normalization modules in SPM12 (http://www.fil.ion.ucl.ac.uk/spm). Using this approach, individual T1-weighted images were segmented into gray, white, and CSF images then non-linearly registered to the existing IXI template of 550 healthy subjects averaged in standard Montreal Neurological Institute space, available with VBM8 (http://dbm.neuro.uni-jena.de/vbm/). Subsequently, gray matter images were scaled by the Jacobian determinant of the transformation to preserve the total amount of signal from each region, and smoothed with an 8mm FWHM Gaussian kernel. The voxel size of processed images was 1.5×1.5×1.5 mm. A gray matter mask for subsequent analyses was created by thresholding the final stage (6th) IXI template at 0.1. Whole-brain voxel-based morphometry maps for each factor were thresholded at voxel-wise p < 0.05, FWE-corrected for multiple comparisons.

As estimates of grey matter volume conflate cortical thickness and surface area, and grey matter volume has a higher correlation with surface area than thickness (14), we did not expect the VBM analyses to recapitulate our surface-based findings. The results of our VBM analyses (Table S2; Figure S10) confirmed our expectations. The paucity of VBM associations across all four factors likely reflects the specificity of our surface-based findings largely to cortical thickness. The biased representation of surface area and cortical thickness in grey matter volume is further reflected in the observation that the Externalizing factor exhibited the highest number of clusters with significantly reduced grey matter volume as total surface area was only correlated with the Externalizing factor in our primary analyses.

As shown in Table S2, we found reduced grey matter volume predominantly within the superior frontal gyrus including orbital and medial regions, thalamus, and cuneus within the occipital cortex in individuals with high *p* factor scores. Our findings are consistent with previous research showing reduced prefrontal grey matter volume within medial and orbitofrontal regions in youth with high *p* factor scores (15). Smaller thalamic volume has been previously identified as a common structural neural correlate in a transdiagnostic VBM meta-analysis (16). Reduced thalamic grey matter volume in individuals with high *p* factor scores is also consistent with our prior research showing structural alterations within a cerebello-thalamo-cortical circuit involved in executive control in young adults with high *p* factor scores (17,18). In these same studies, we also found reduced grey matter volume within the visual association cortex in those with high *p* factor scores, which specifically overlaps with our current finding of reduced grey matter volume in the cuneus region of occipital cortex.

Prior VBM meta-analyses have reported smaller grey matter volume of the dorsal anterior cingulate cortex (dACC) and insula across diagnostic categories (16,19). Our VBM analyses did not reveal similar associations with the *p* factor. However, our methods for examining phenotypic commonalities across diagnostic categories are different from these meta-analyses, which employed case-control designs and tested for convergence across diagnostic groups vs. healthy controls. The *p* factor captures shared variation across diagnostic categories essentially measuring severity and comorbidity of psychiatric disorders continuously, which is substantially different from the approach of collapsing across diagnostic categories. Thus, we would not necessarily expect our VBM findings to completely match those of previous meta-analyses. Differences in age also could be a factor contributing to mismatches between our current and previous VBM findings. For example, recent research by Kaczkurkin et al. (20) showed reduced global grey matter volume and cortical thinning in only two of 18 cortical networks in youth with high *p* factor scores (aged 8-21). Our results may differ from those of Kaczkurkin et al. (20) due to the difference in ages between the two samples (youth vs. midlife adults aged 45), suggesting possible developmental changes in grey matter volume and cortical thickness patterns in those with high general psychopathology throughout the life course.

3.	APA. American Psychological Association: Diagnostic and Statistical Manual of Mental Disorders (Revised Third ed.). Washington, DC: American Psychiatric Association; 1987.

4.	APA. American Psychological Association: Diagnostic and Statistical Manual of Mental Disorders (Fourth ed.). Washington, DC: American Psychiatric Association; 1994.

5.	APA. American Psychological Association: Diagnostic and Statistical Manual of Mental Disorders (Fifth ed.). Washington, DC: American Psychiatric Association; 2013.

6.	Caspi A, Houts RM, Belsky DW, Goldman-Mellor SJ, Harrington H, Israel S, et al. The p factor: One general psychopathology factor in the structure of psychiatric disorders? Clin Psychol Sci J Assoc Psychol Sci. 2014 Mar;2(2):119–37.

7.	Heatherton TF, Kozlowski LT, Frecker RC, Fagerström KO. The Fagerström Test for Nicotine Dependence: a revision of the Fagerström Tolerance Questionnaire. Br J Addict. 1991 Sep;86(9):1119–27.

8.	Caspi A, Moffitt TE. All for one and one for all: mental disorders in one dimension. Am J Psychiatry. 2018 Apr 6;175(9):831–44.

9.	Muthen LK, Muthen BO. Mplus User’s Guide. (Eight ed.). Los Angeles, CA: Muthen & Muthen; 1998.

10.	Asparouhov T, Muthen B. Weighted Least Squares Estimation with Missing Data. Available from: Retrieved from http://www.statmodel.com/download/GstrucMissingRevision.pdf

11.	Bollen KA, Curran PJ. Latent Curve Models A Structural Equation Perspective Introduction. [Internet]. 2006. 15 p. Available from: Retrieved from <Go to ISI>://WOS:000297795700002.

**Table S1.** Standardized factor loadings for models of the structure of psychopathology.

|  |  | Model A: Correlated Factors | | |  | Model B': Bifactor Model | | |
| --- | --- | --- | --- | --- | --- | --- | --- | --- |
| Model Fit Statistics | |  |  |  |  |  |  |  |
|  | Chi-Square (WLSMV) | 4082.230 | | |  | 3695.364 | | |
|  | Degrees of Freedom | 2465 | | |  | 2457 | | |
|  | Comparative Fit Index | 0.933 | | |  | 0.949 | | |
|  | Tucker-Lewis Index | 0.929 | | |  | 0.945 | | |
|  | RMSEA [90% CI] | 0.026 [0.024, 0.027] | | |  | 0.022 [0.021, 0.024] | | |
|  |  | Externalizing | Internalizing | Thought Disorder |  | *p* factor | Externalizing | Internalizing |
| Standardized factor loadings | | |  |  |  |  |  |  |
|  | ADHD | 0.567 |  |  |  | 0.595 | 0.121 |  |
|  | Alcohol | 0.651 |  |  |  | 0.300 | 0.622 |  |
|  | Cannabis | 0.831 |  |  |  | 0.369 | 0.850 |  |
|  | Hard drugs | 0.845 |  |  |  | 0.466 | 0.694 |  |
|  | Tobacco | 0.675 |  |  |  | 0.450 | 0.468 |  |
|  | Conduct disorder | 0.888 |  |  |  | 0.504 | 0.714 |  |
|  | Major depression |  | 0.968 |  |  | 0.768 |  | 0.587 |
|  | Generalized anxiety |  | 0.892 |  |  | 0.686 |  | 0.642 |
|  | Fears/phobias |  | 0.717 |  |  | 0.582 |  | 0.424 |
|  | Eating disorder |  | 0.499 |  |  | 0.377 |  | 0.374 |
|  | PTSD |  | 0.994 |  |  | 0.860 |  | 0.351 |
|  | OCD |  |  | 0.739 |  | 0.772 |  |  |
|  | Mania |  |  | 0.955 |  | 0.976 |  |  |
|  | Schizophrenia |  |  | 0.838 |  | 0.865 |  |  |
| Factor Correlations | |  |  |  |  |  |  |  |
|  | Externalizing |  | 0.420 | 0.622 |  |  |  |  |
|  | Internalizing |  |  | 0.847 |  |  |  |  |

**Figure S1.** **Attrition analyses show scanned study members are representative of the entire cohort on measures of (A) childhood IQ, (B) SES, and (C) p factor scores.**

**(A)**

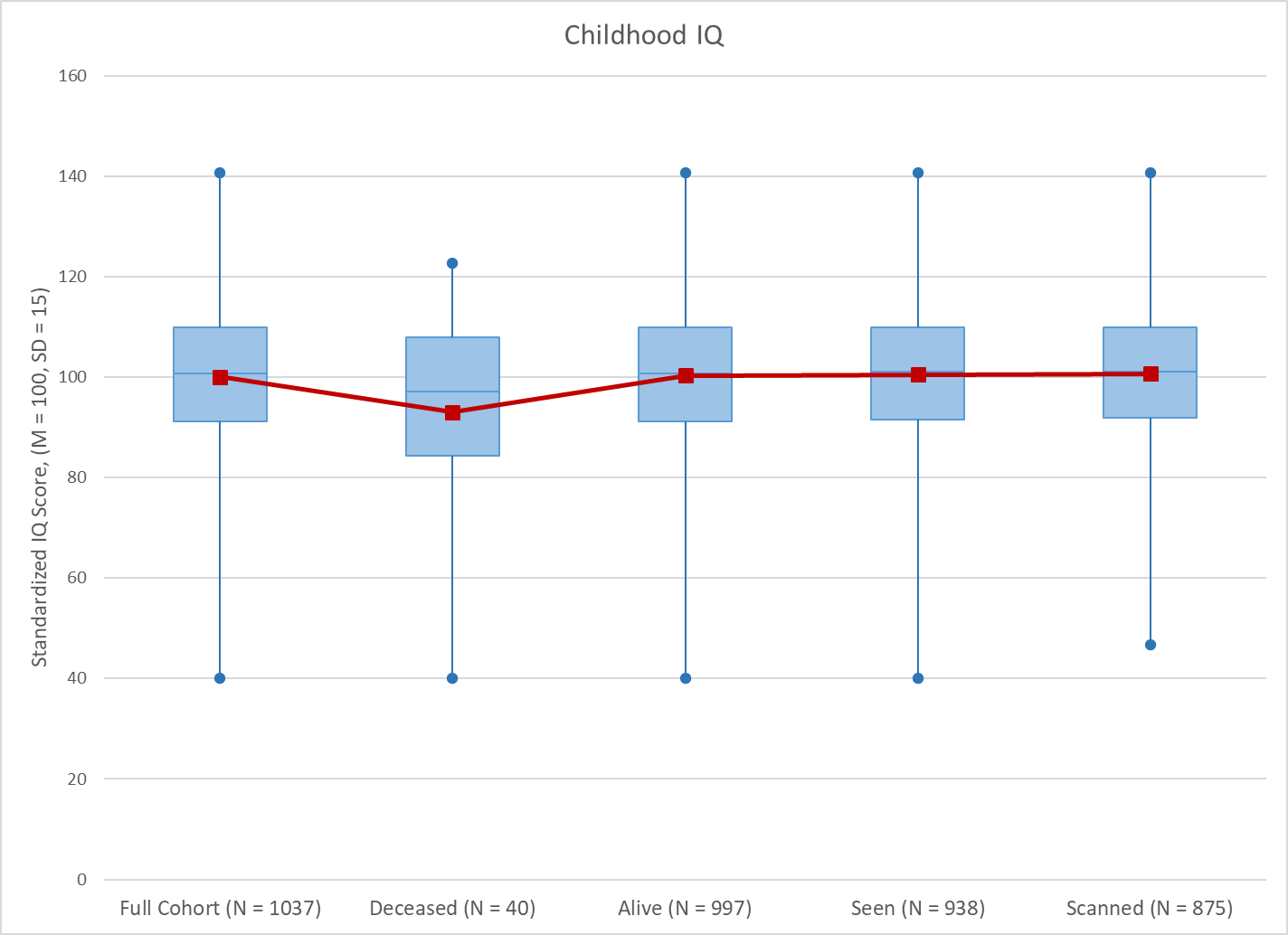

Those who were deceased by the Phase 45 data collection had significantly lower childhood IQ’s than those who were still alive (t = 2.09, p = 0.04).

**(B)**

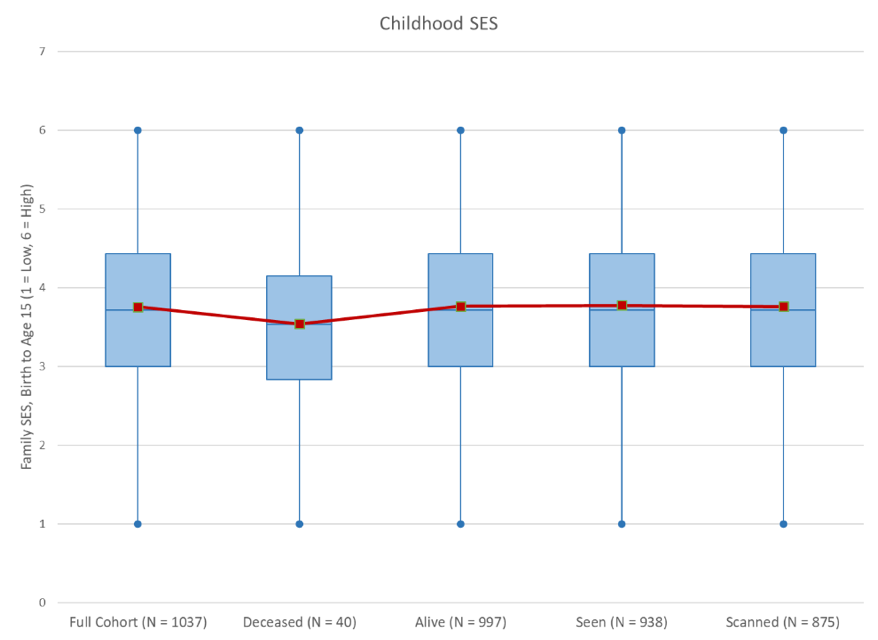

No significant differences were found between the full cohort, those deceased, those alive, those seen at Phase 45 or those scanned at Phase 45 on childhood SES.

(**C)**

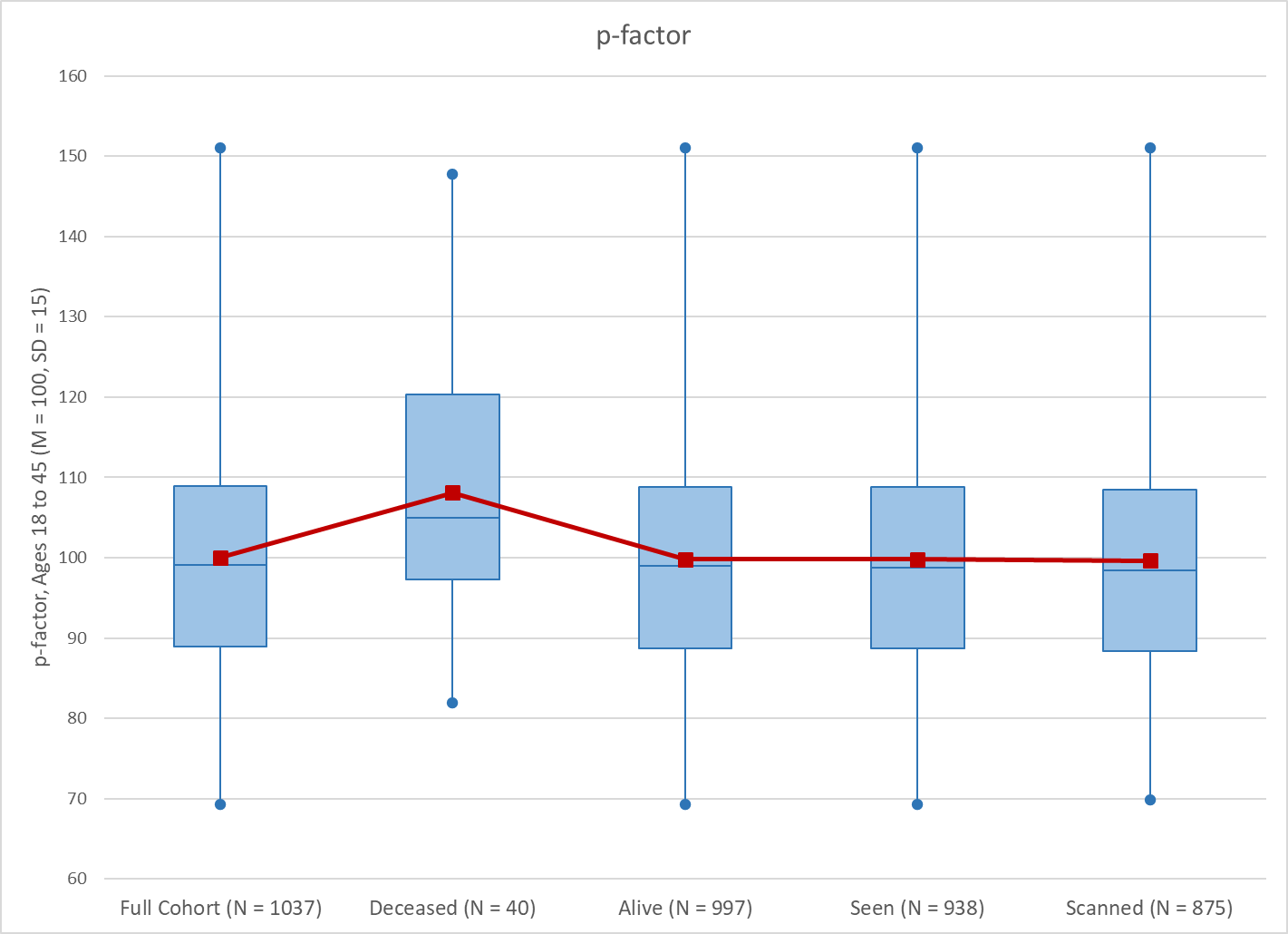

No significant differences in *p* factor scores were found between the full cohort, those still alive, those seen at Phase 45 or those scanned at Phase 45. Those who were deceased by the Phase 45 data collection had significantly higher *p* factor scores than those who were still alive (t = 2.86, p = 0.004).

**Figure S2.** **Structure of mental-disorder data collected in the Dunedin Study.**

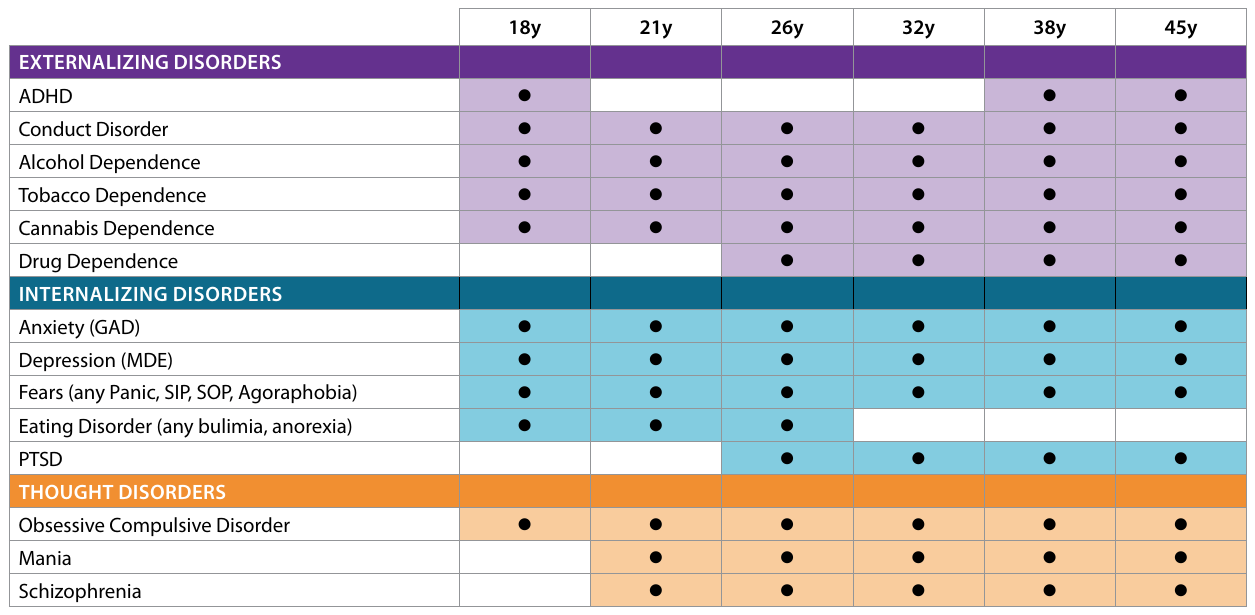

The chart shows the age at which each disorder was assessed. Although each disorder was not assessed at every age, each disorder was assessed on at least three occasions.

**Figure S3**. **Confirmatory factor analysis models of the structure of psychopathology.**

**Model A**

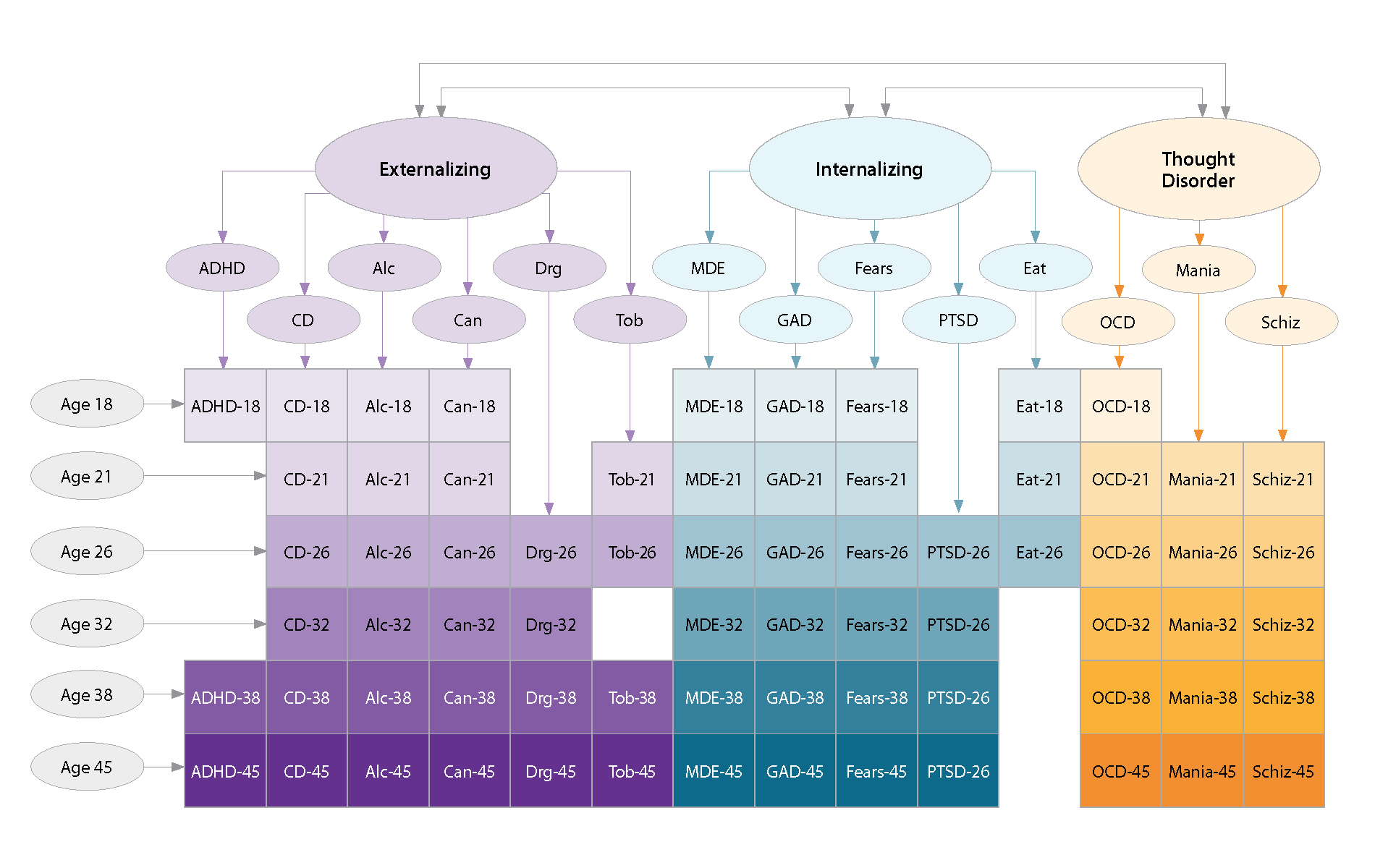

**Model A** is the correlated-factors model. Using this model, we tested the hypothesis that there are latent trait factors, each of which influences a subset of the diagnostic symptoms. We tested three factors representing Externalizing (with loadings from ADHD, conduct disorder, alcohol dependence, cannabis dependence, drug dependence and tobacco dependence), Internalizing (with loadings from MDE, GAD, fears/phobias, PTSD, and eating disorders), and Thought Disorder (with loadings from OCD, mania, and schizophrenia).

**Model B**

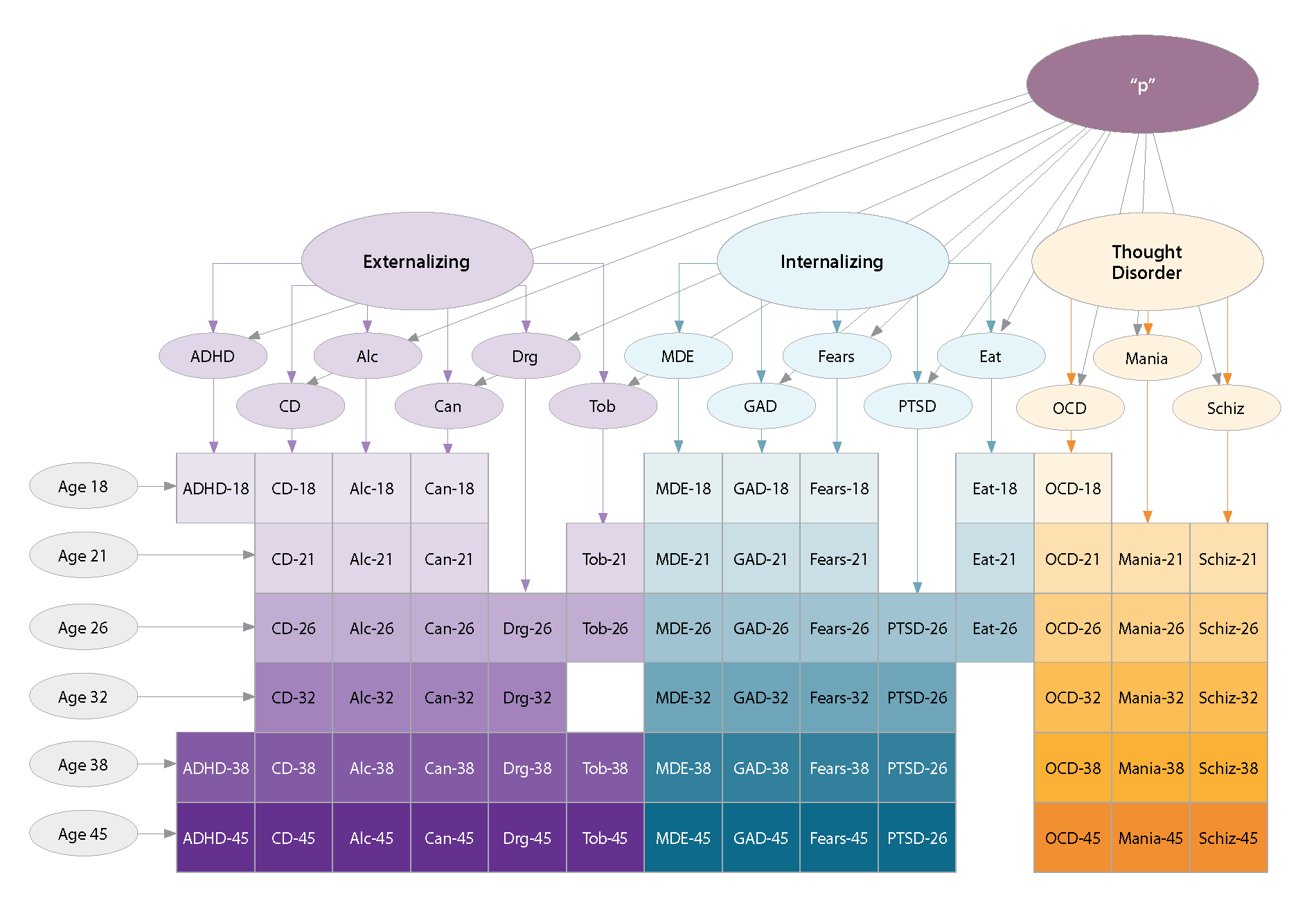

**Model B** is the hierarchical or bifactor model. Using this model, we tested the hypothesis that the symptom measures reflect both General Psychopathology and three narrower styles of psychopathology. General Psychopathology (labeled *p* in Model B) is represented by a factor that directly influences all of the diagnostic symptom factors. In addition, styles of psychopathology are represented by three factors, each of which influences a smaller subset of the symptom items. For example, alcohol symptoms load jointly on the General Psychopathology factor and on the Externalizing style factor. The specific factors represent symptoms of Externalizing, Internalizing, and Thought Disorder that are independent of General Psychopathology.

**Model B’**

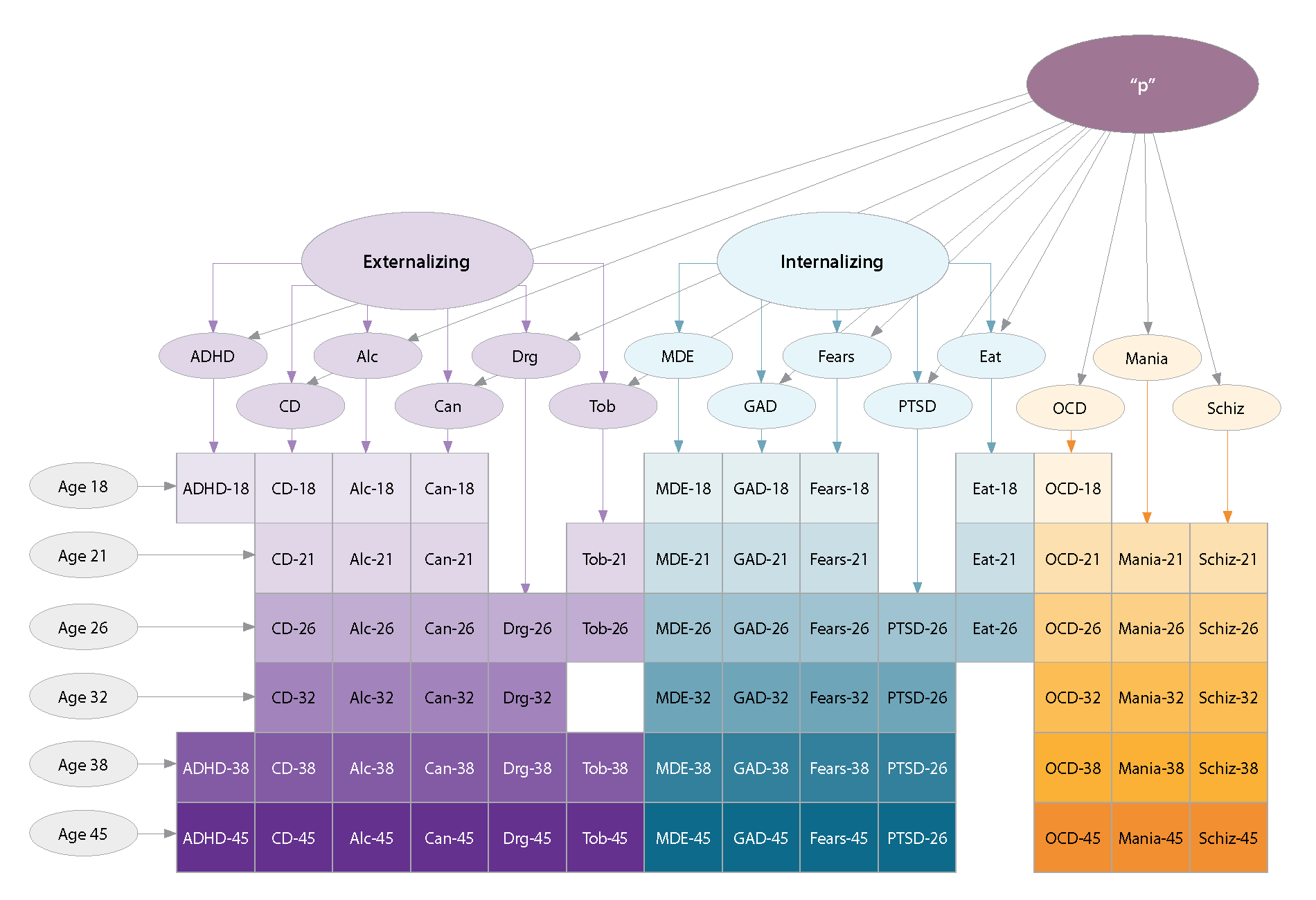

Model B had a Heywood case, an estimated variance that was negative for one of the lower-order disorder/symptom factors (specifically, mania), suggesting this was not a valid model. Inspection of the results revealed the source of the convergence problem. Specifically, the Thought Disorder factor was subsumed in *p*; that is, in the hierarchical model, symptoms of OCD, mania, and schizophrenia loaded very highly on *p*, but unlike symptoms of Externalizing and Internalizing, they could not form a separate Thought Disorder factor independently of *p*. We respecified the model accordingly, depicted in **Model B′**.

**Figure S4.** **Plot of the positive correlation between the variety and persistence of mental disorders and *p* factor scores in the Dunedin Study.**

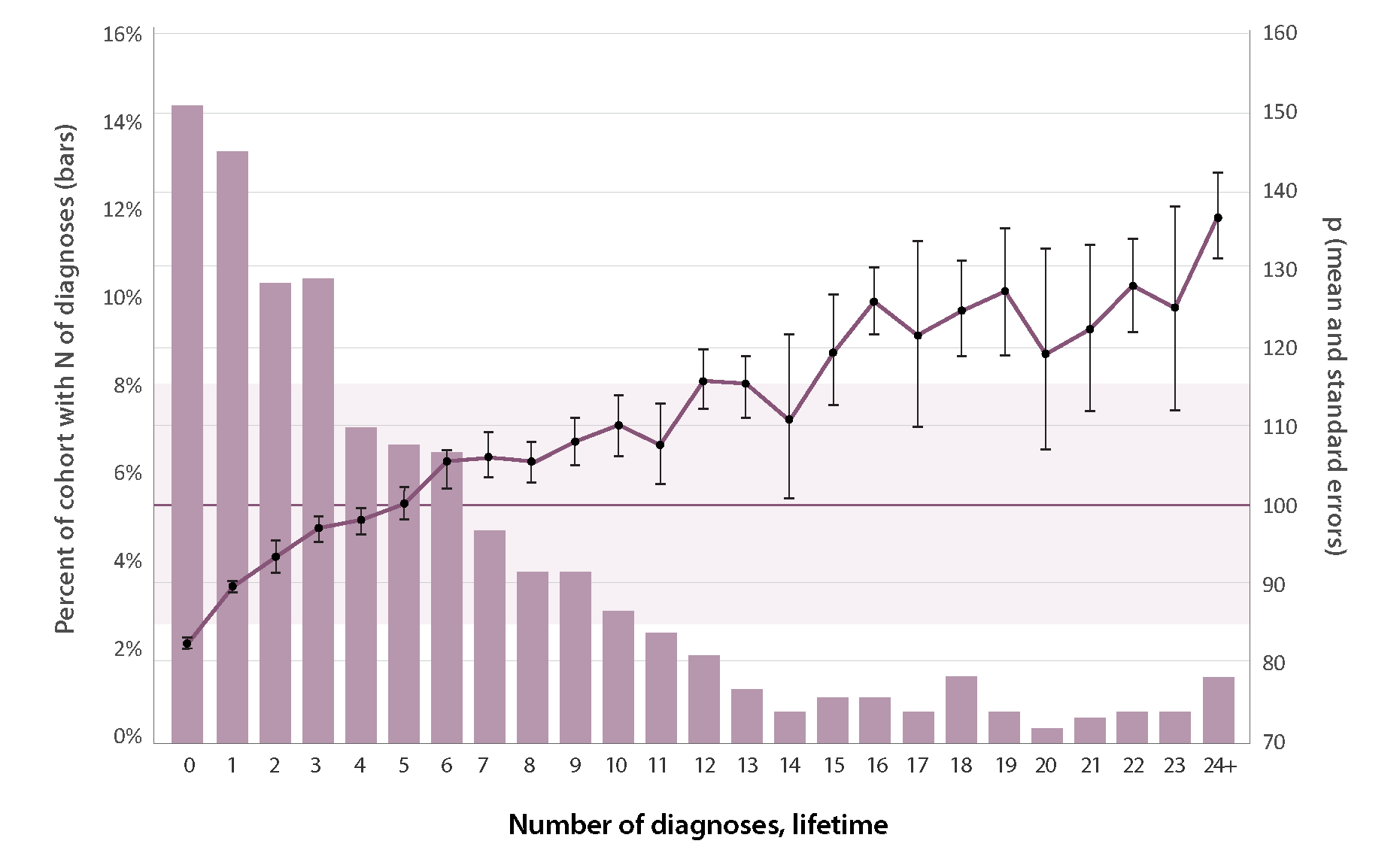

The *p* factor captures how Study members differ from each other in the variety and persistence of many different kinds of disorders over the life course. Study members with higher *p* scores experienced a greater number of mental disorders from adolescence to midlife (r=.76). Shaded area represents ± 1 SD for *p*.

**Figure S5.** **Covariate analyses of associations between factor scores and global cortical thickness.** We conducted analyses with the addition of several covariates to our initial covariate of sex focusing on global cortical thickness. Specifically, we modeled if Study members reported taking any psychoactive medication at age 45 or if they were diagnosed with a chronic medical disease as well as their childhood SES. We also modeled a dimensional measure of Image Quality – the Euler number – derived from FreeSurfer. As illustred below, these analyses revealed that the association between p factor scores and global cortical thickness were robust to all of these covariates.

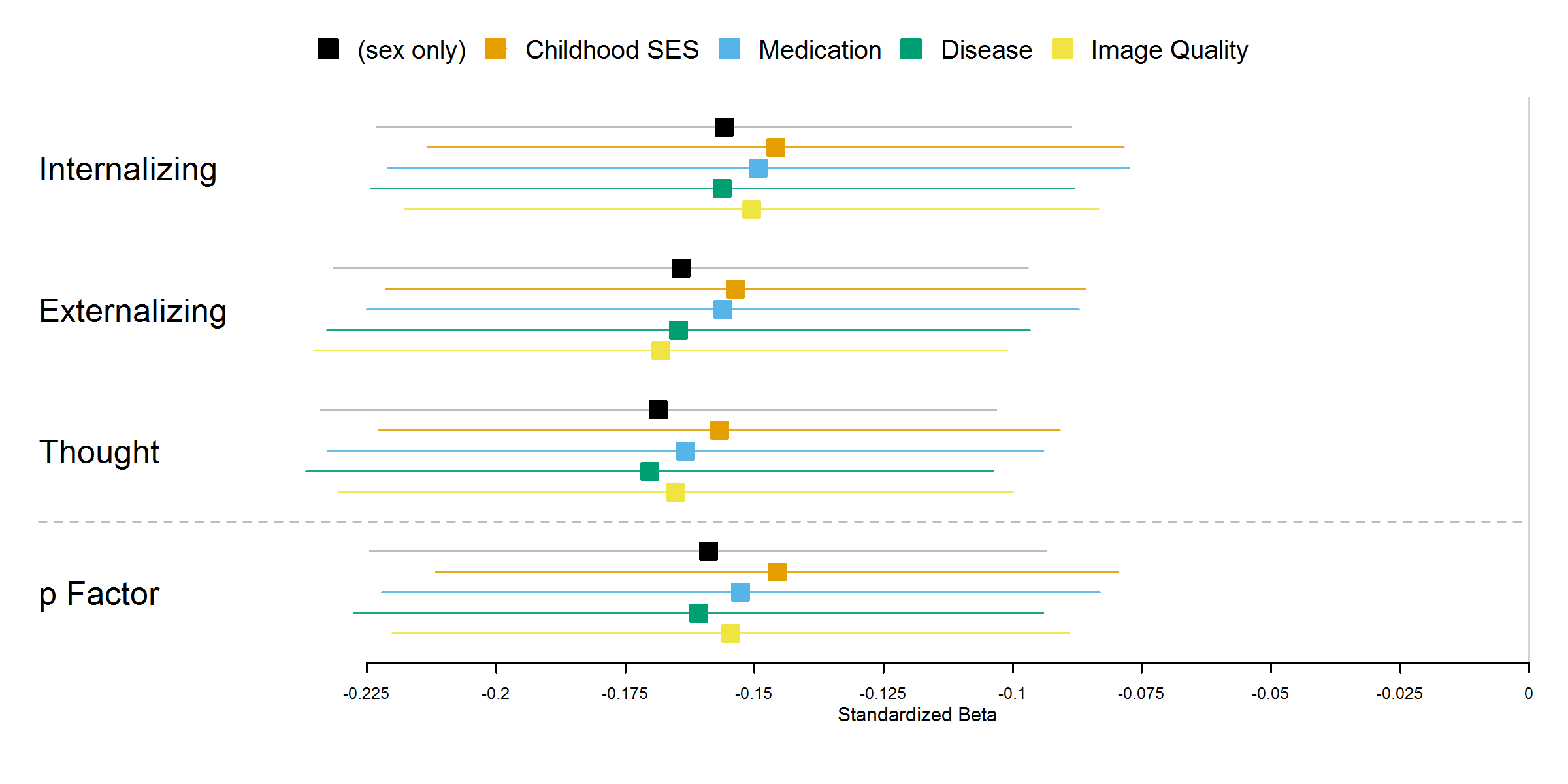

**Figure S6. Case-Control Analyses of Global Cortical Thickness.** Consistent with our factor-based analyses, all of these case-control analyses revealed decreased global cortical thickness as a feature of diagnosis.

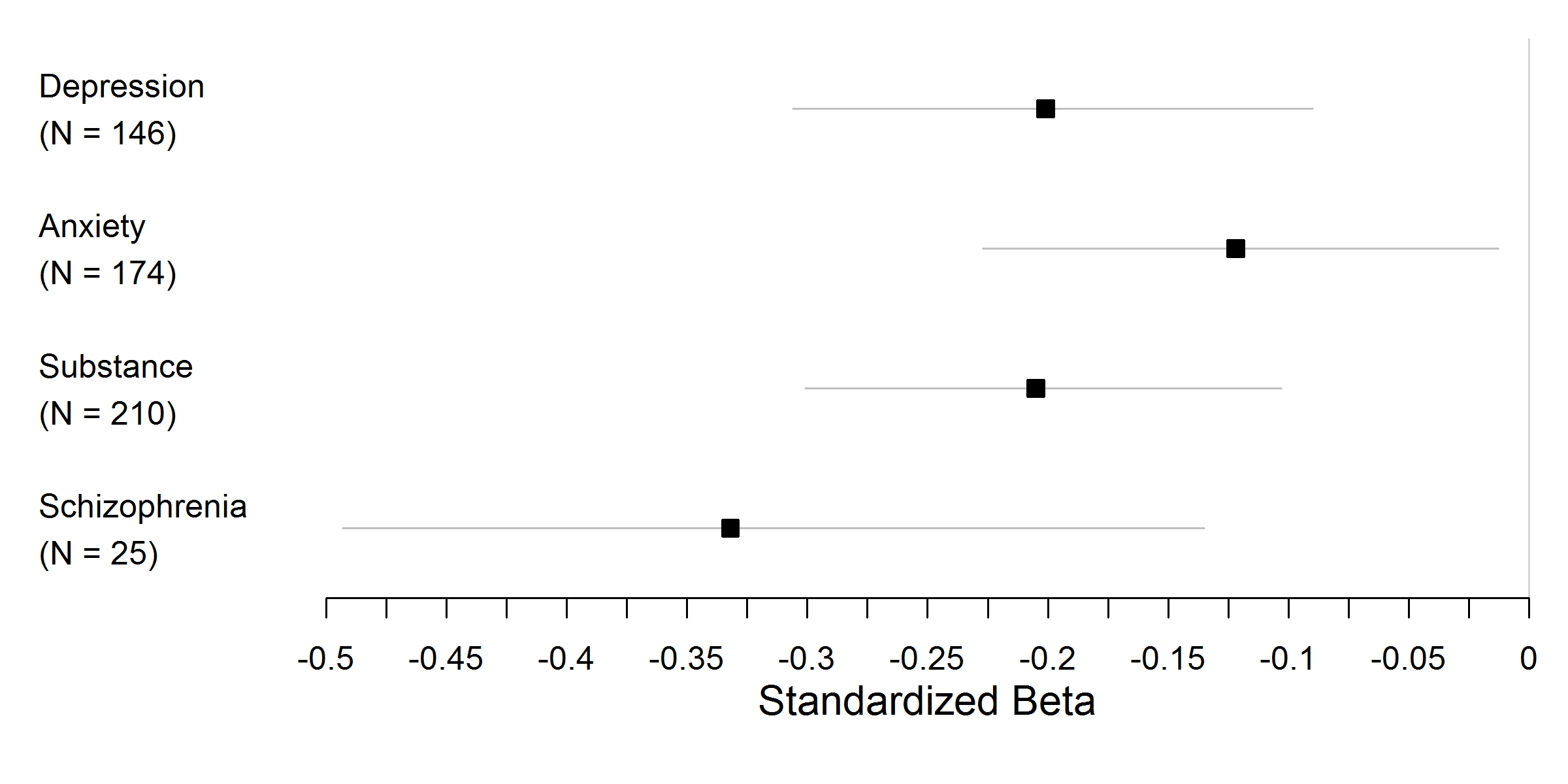

**Figure S7.** **Forest plots depicting the effect sizes (standardized ßs) and 95% confidence intervals of associations between each of the four factor scores and cortical thickness for all 360 parcels, divided into left and right hemisphere columns.** INT: Internalizing; EXT: Externalizing; THT: Thought Disorder; *p*: General Psychopathology.

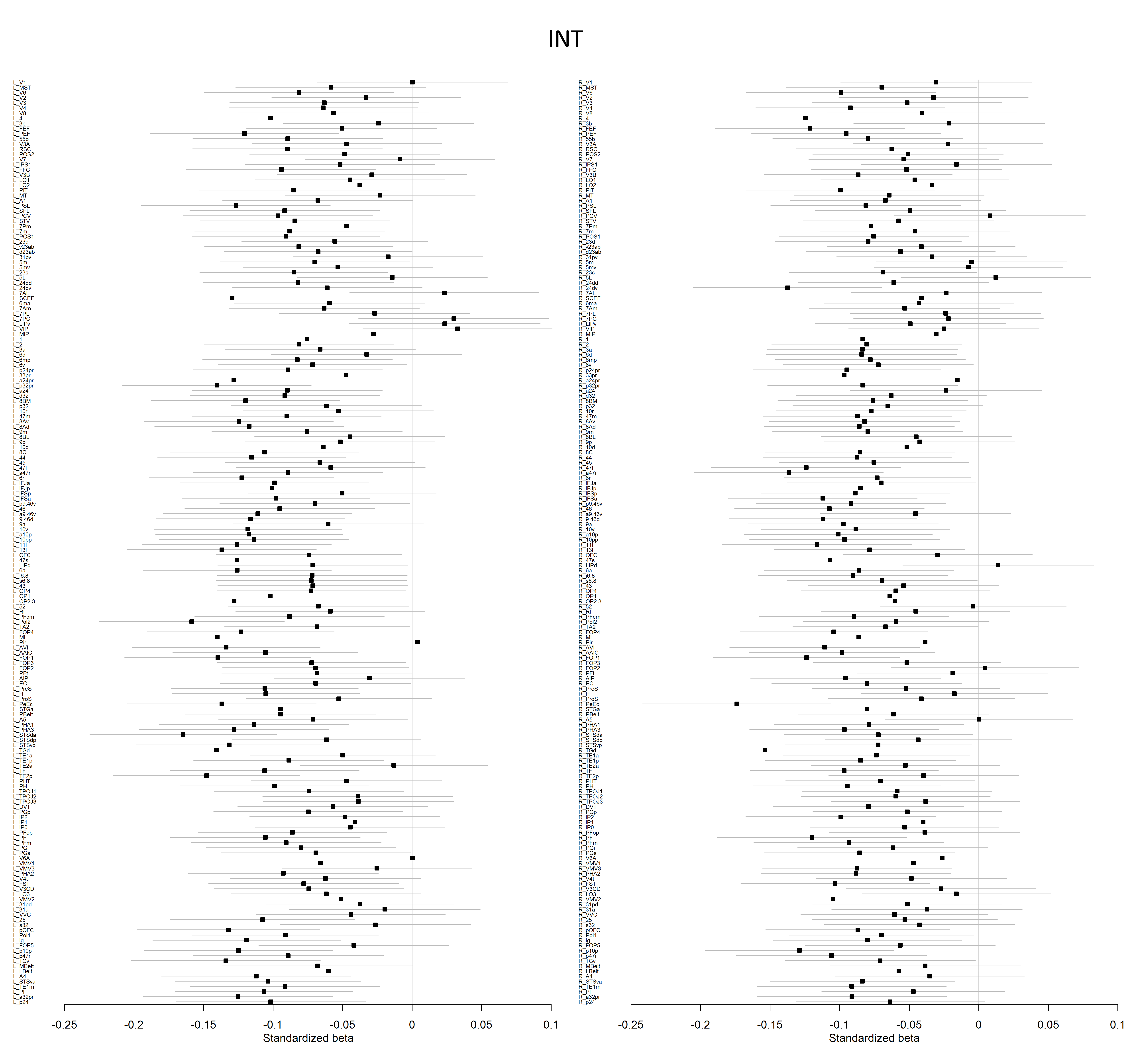

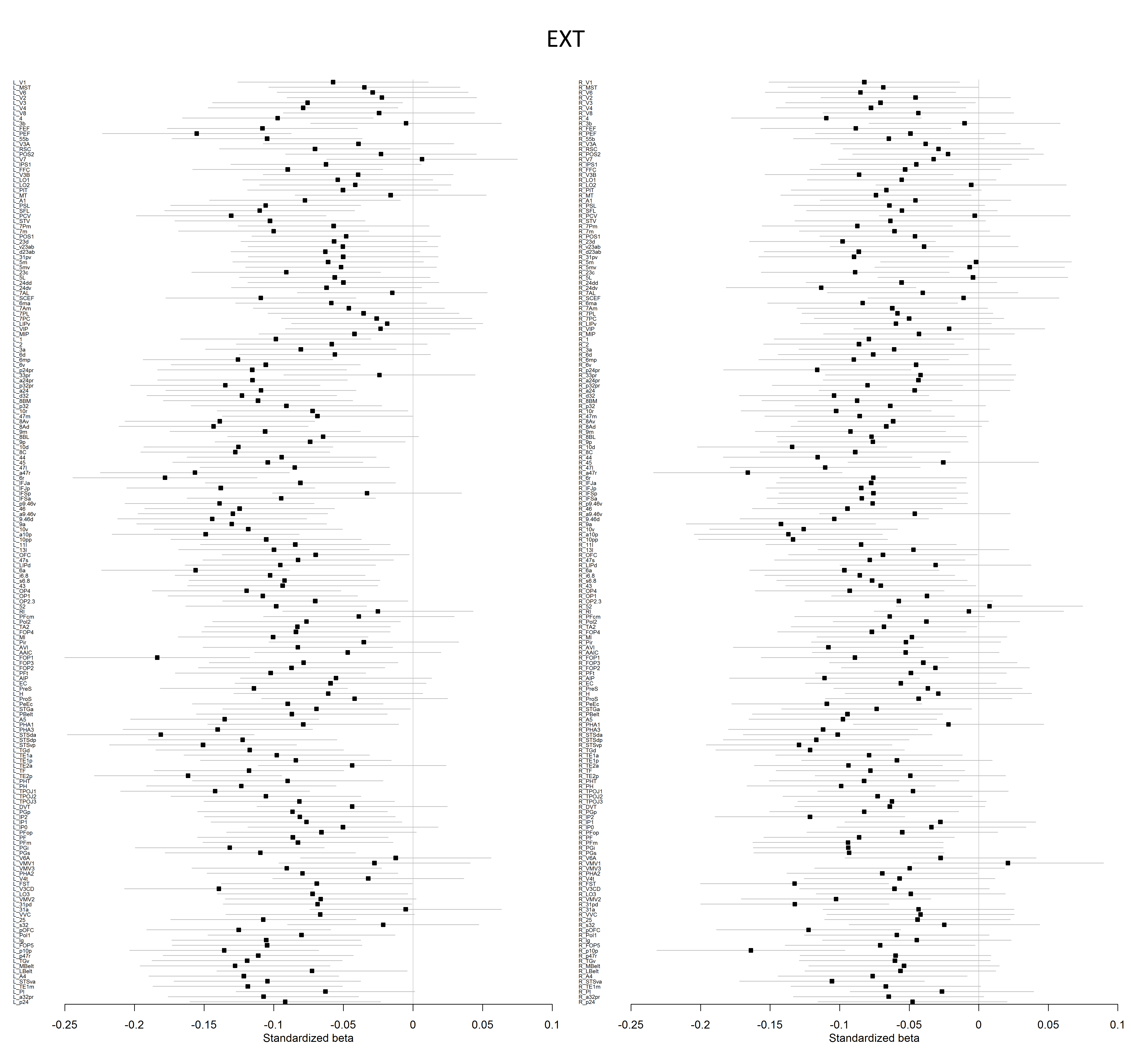

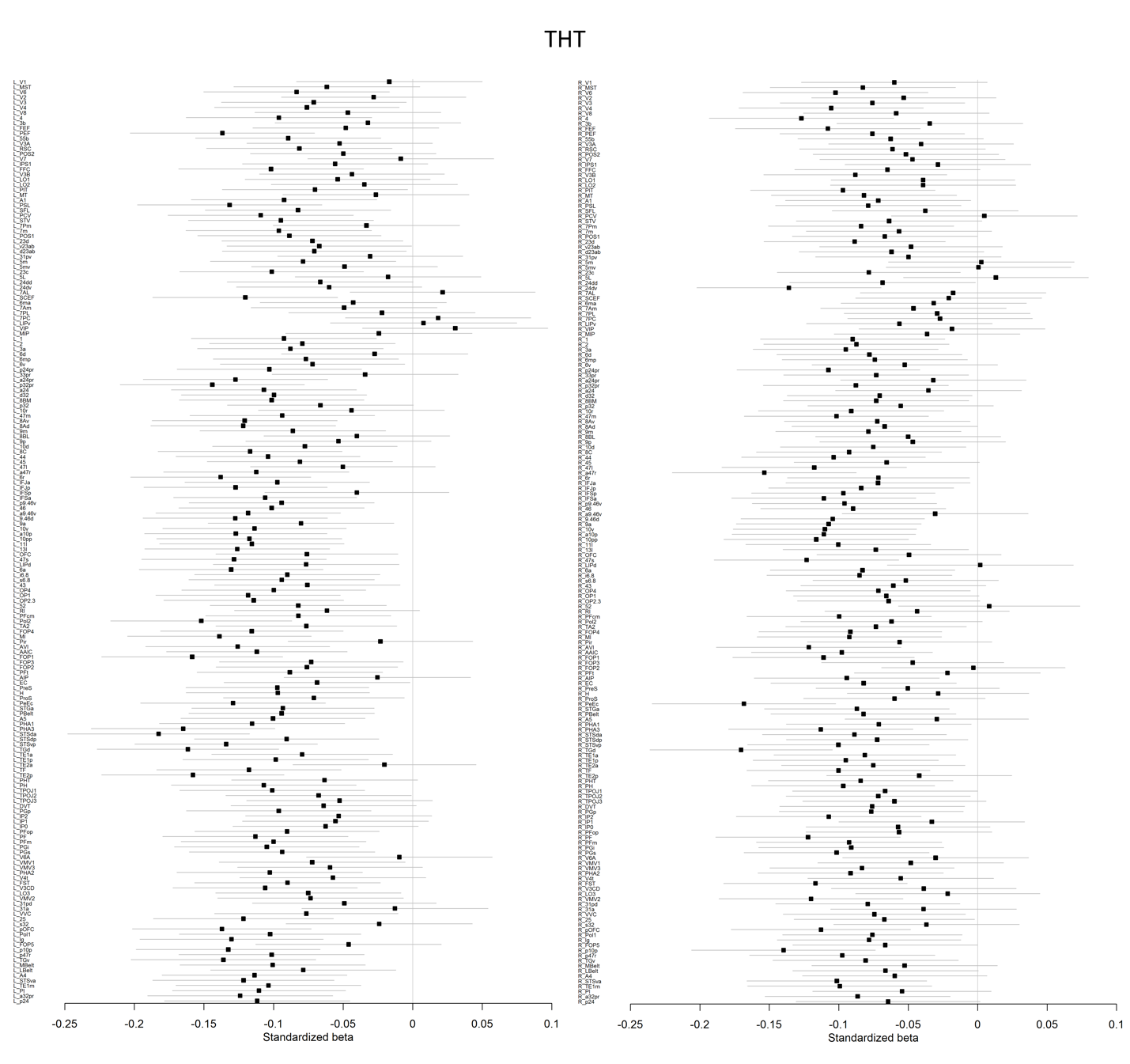

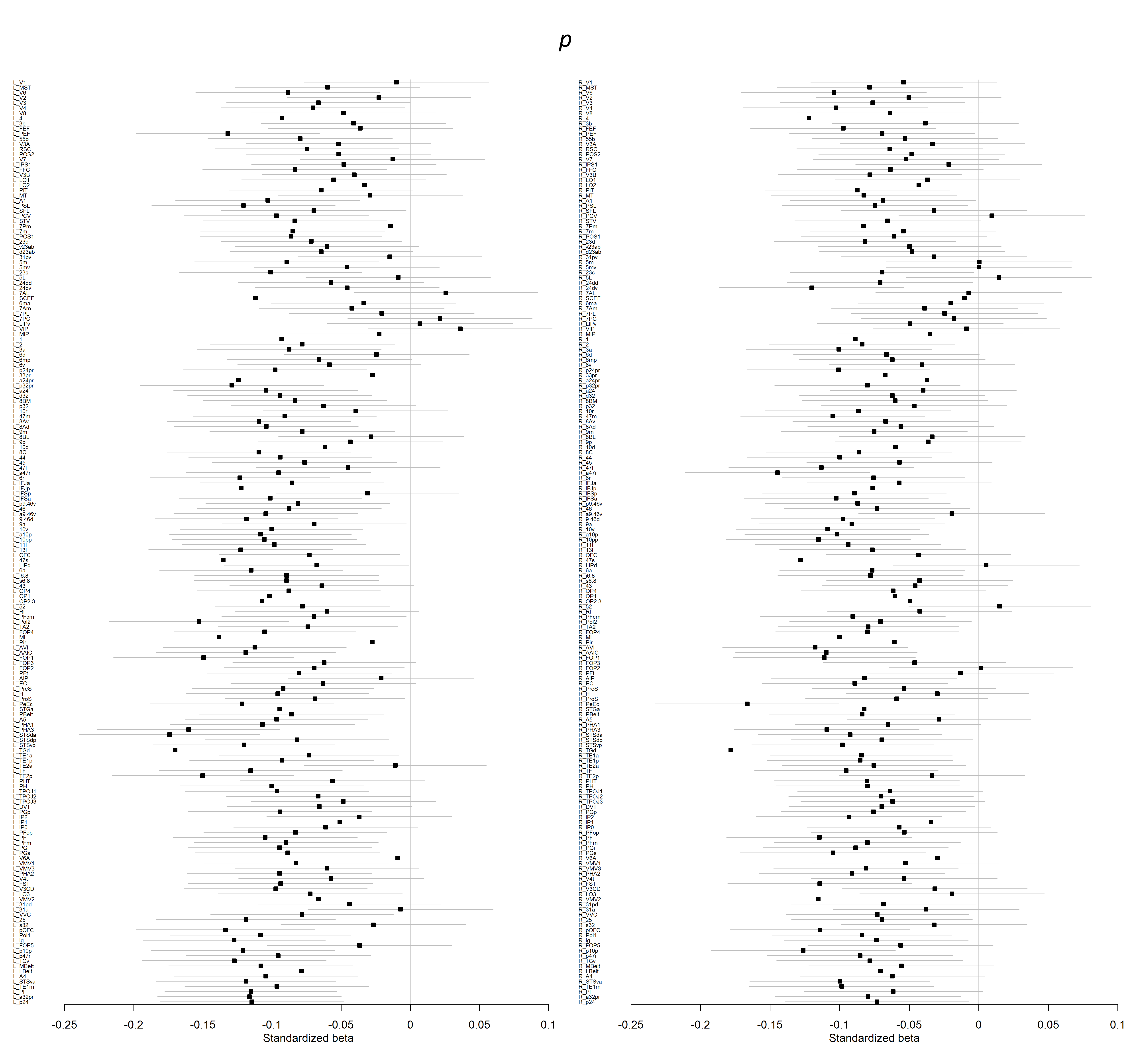

**Figure S8.** **Mutlivariate analysis of unique associations between parcel-wise cortical thickness and scores on the three broad diagnostic families of disorder derived from the correlated factors model.** When partialling out the variance associated with the other factors, a significant association with reduced cortical thickness was only observed between two parcels and scores on the Externalizing factor. There were no unique associations with scores on the Internalizing or Thought Disoder factors. Color bar reflects effect sizes (standardized *β*s).

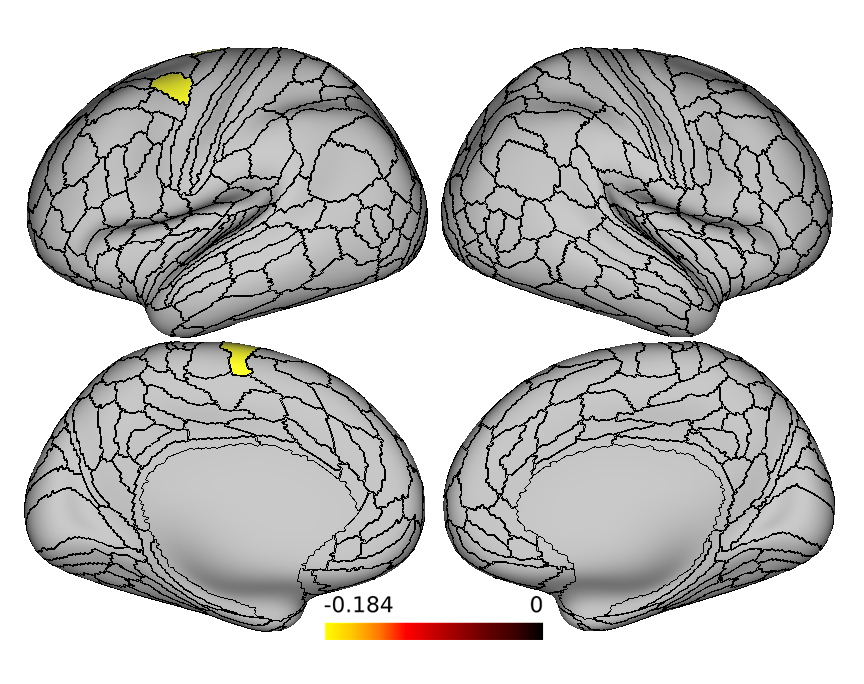

**Figure S9.** **Forest plots depicting the effect sizes (standardized ßs) and 95% confidence intervals of associations between each of the four factor scores and surface area for all 360 parcels, divided into left and right hemisphere columns.** INT: Internalizing; EXT: Externalizing; THT: Thought Disorder; *p*: General Psychopathology.

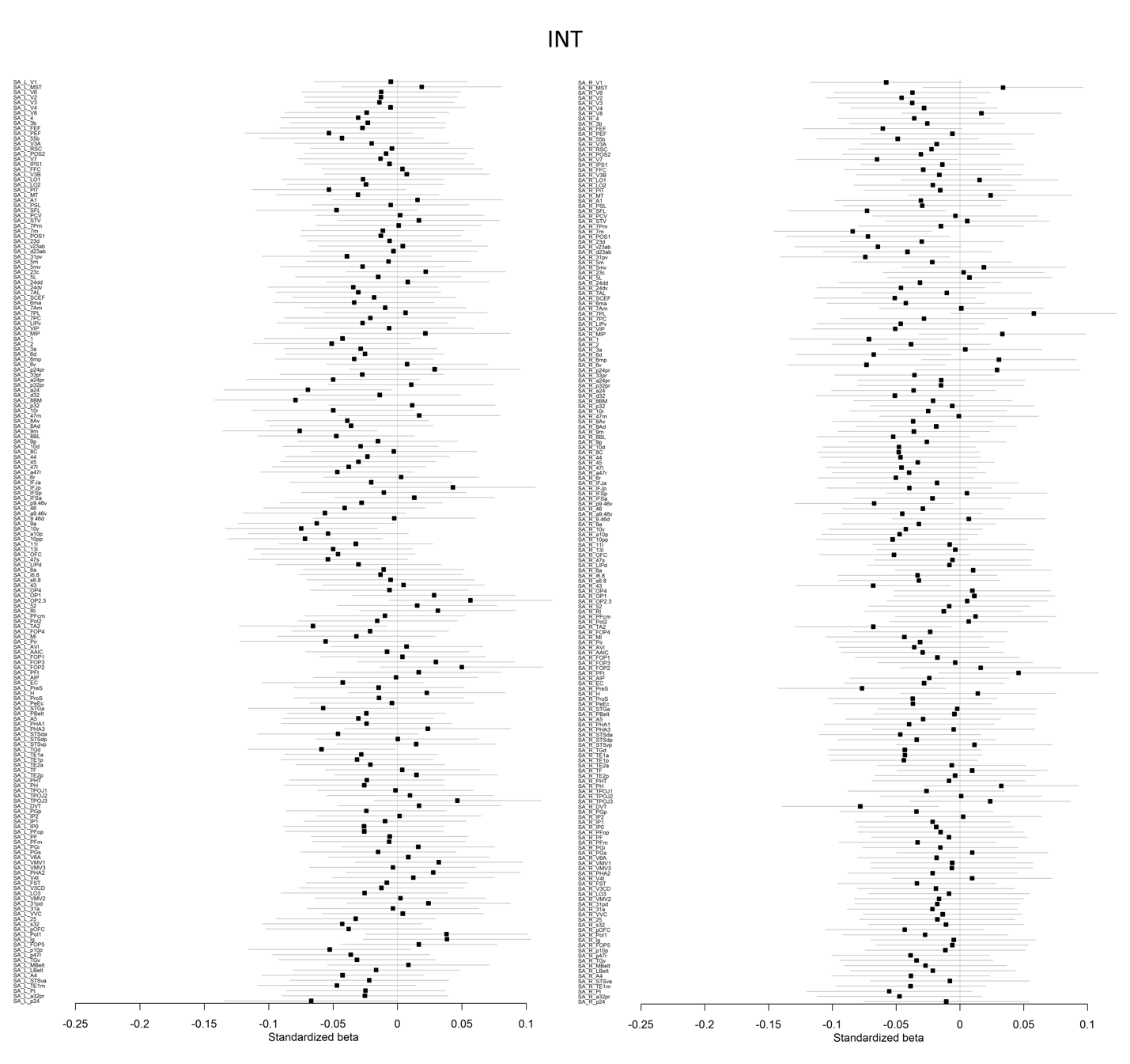

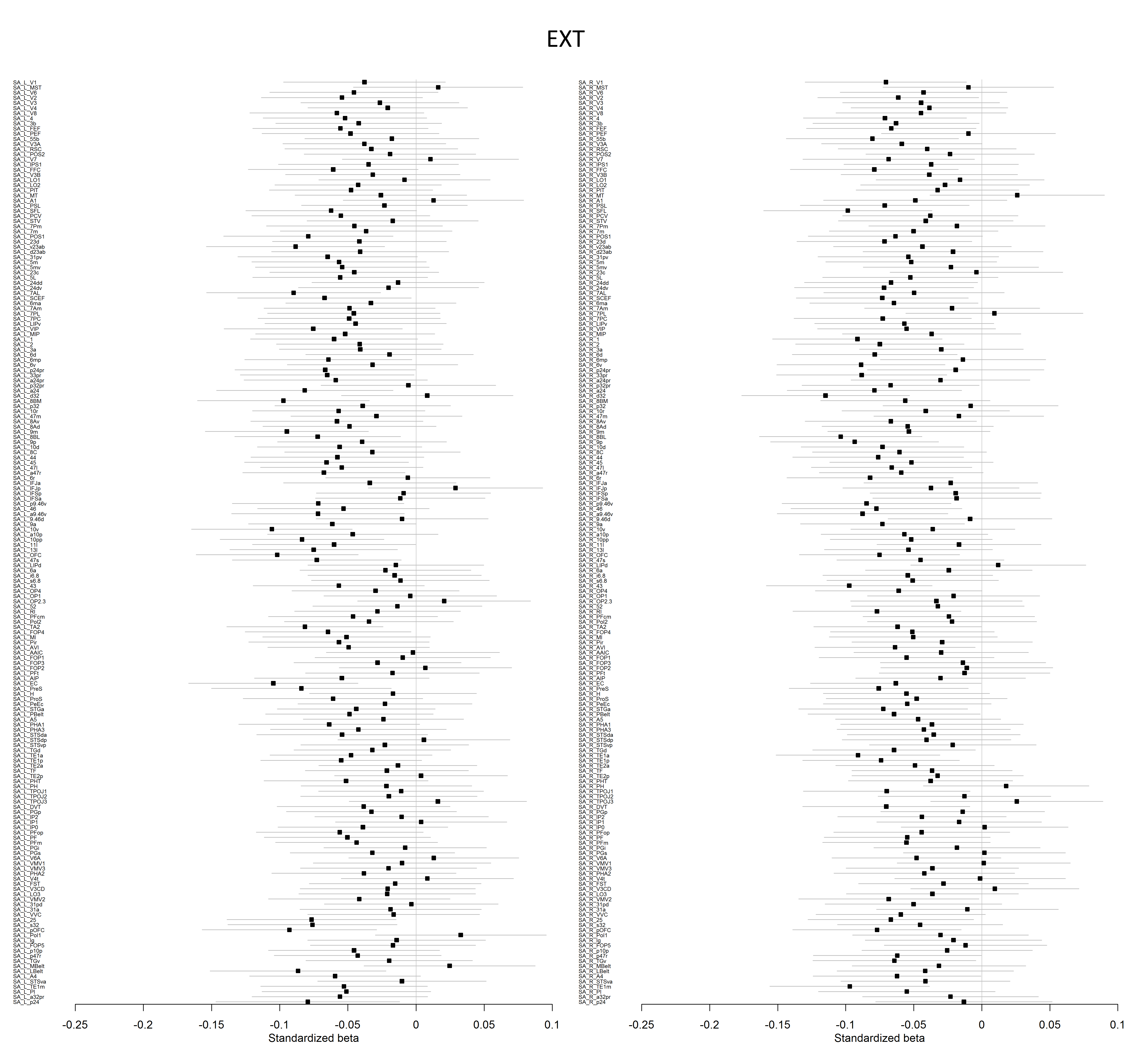

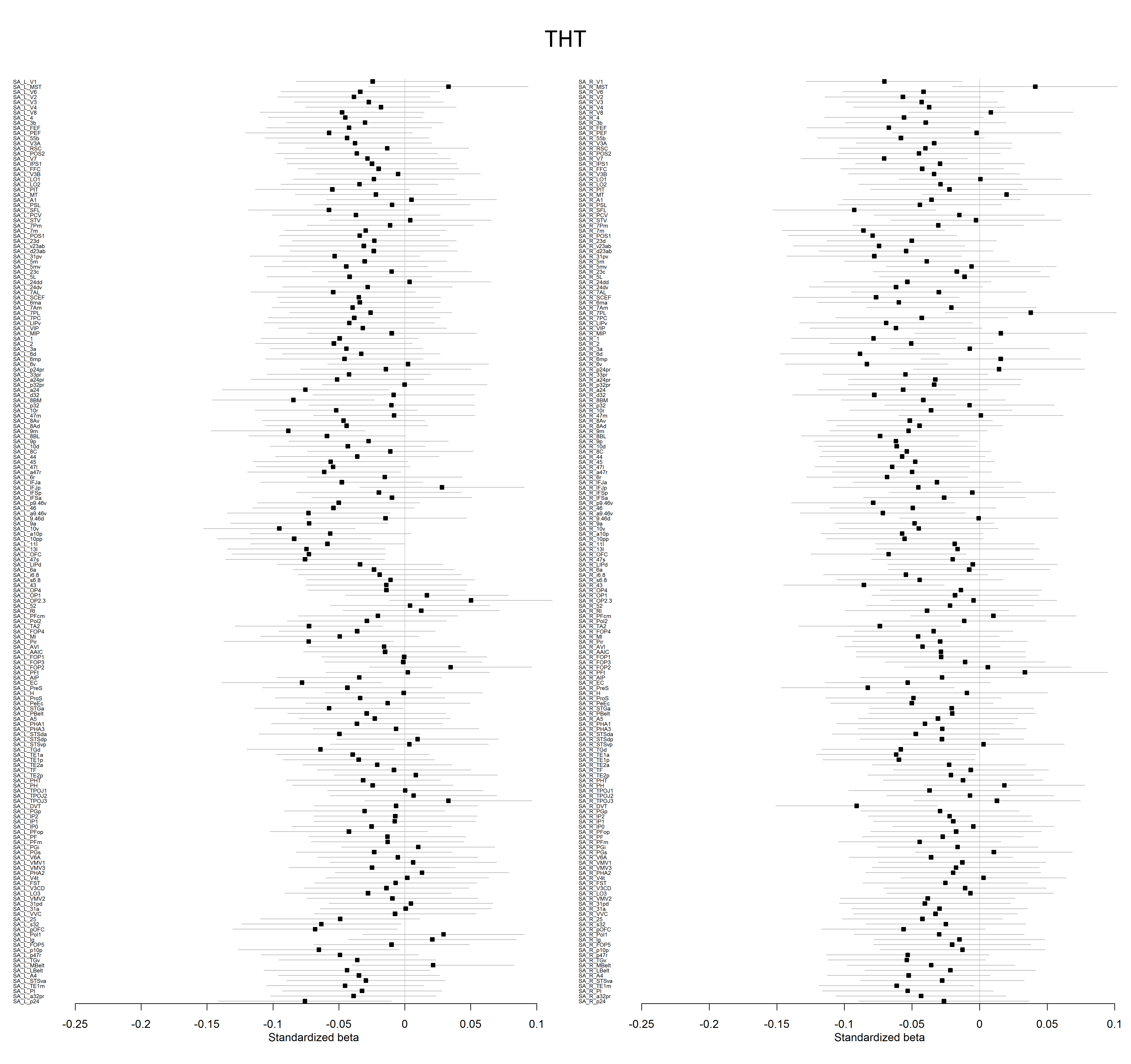

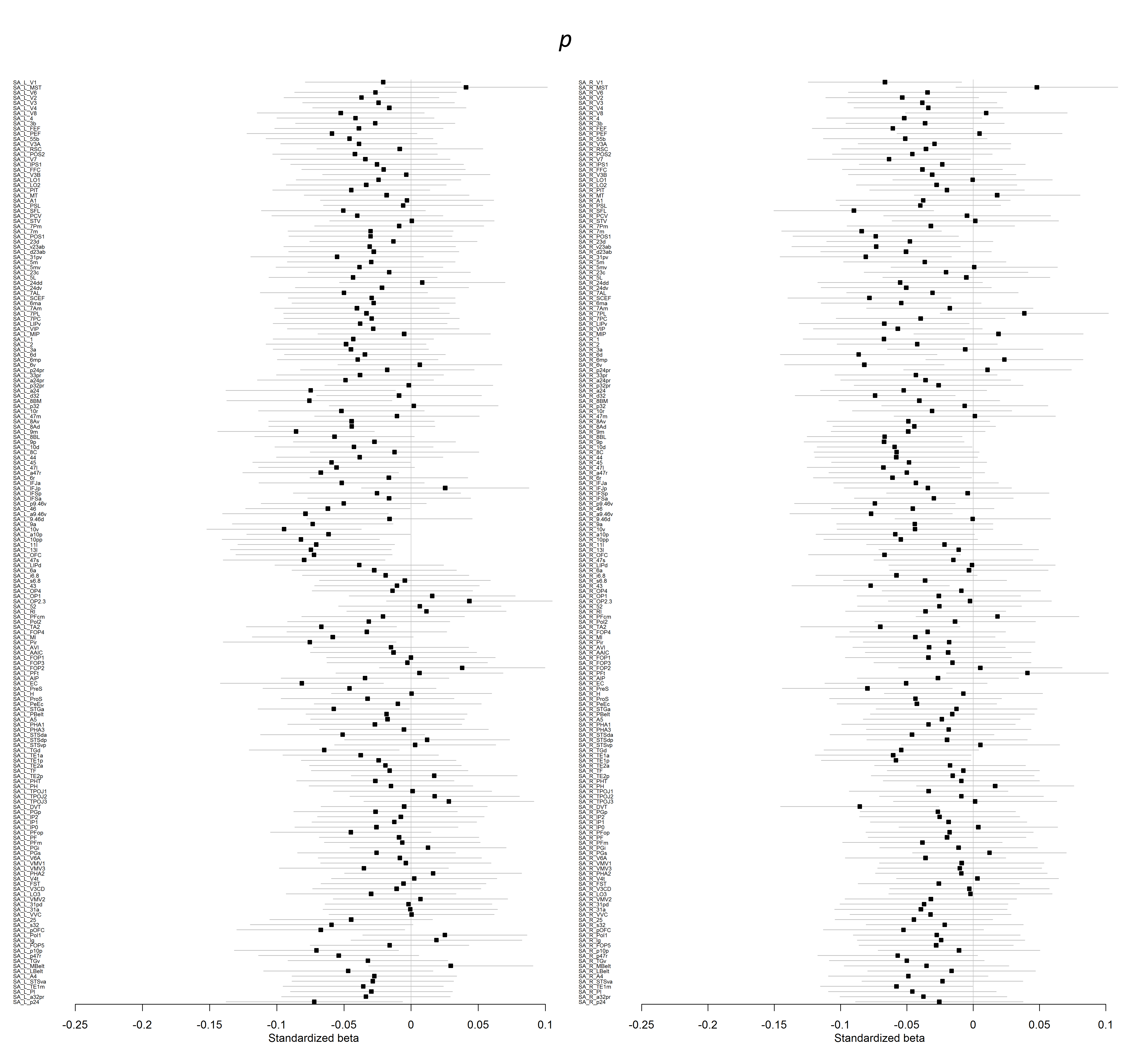

**Figure S10. Whole-brain voxel-wise associations with grey matter volume for each of the four factor scores.** Whole-brain voxel-based morphometry maps for each factor were thresholded at voxel-wise p < 0.05, FWE-corrected for multiple comparisons.

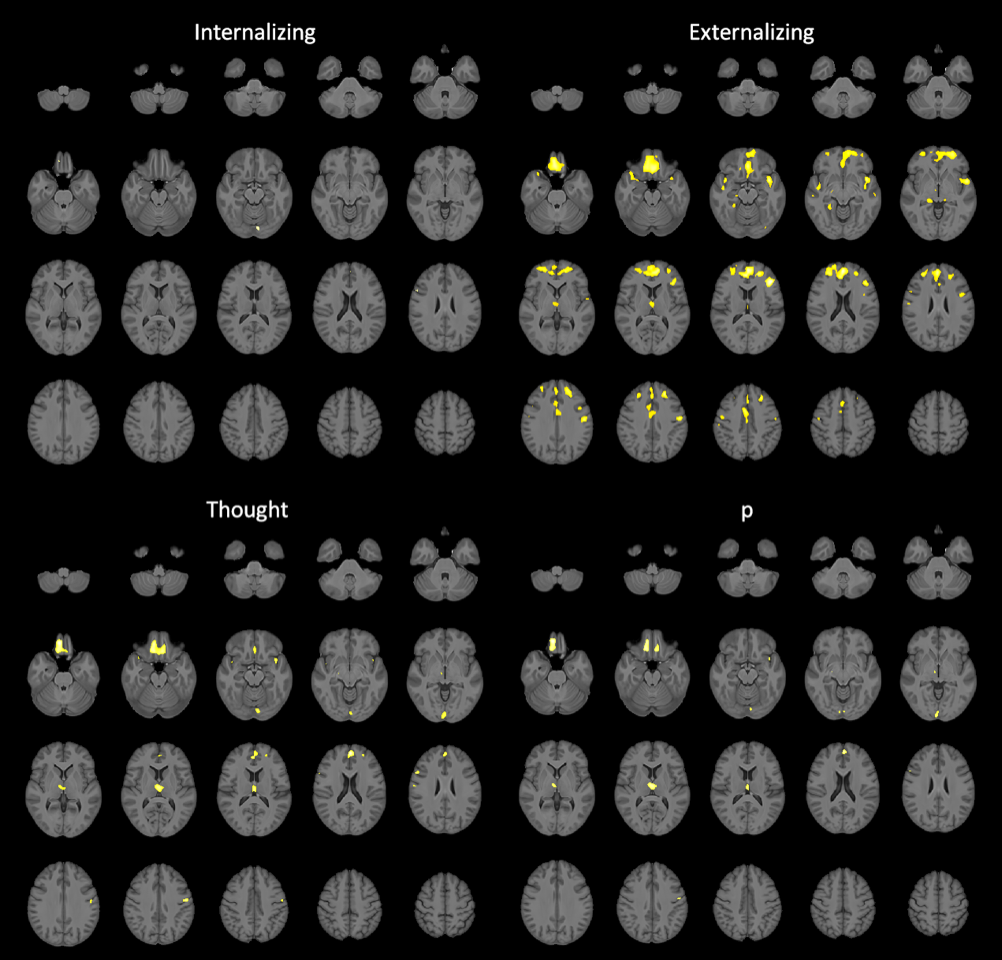

**Table S2.** Summary statistics and anatomical labels for significant clusters in the VBM analyses of grey matter volume correlates of the four factor scores depicted in Figure S10.

| ***Cluster*** | | ***Peak*** | | | |
| --- | --- | --- | --- | --- | --- |
| **p(FWE-corr)** | **k** | **p(FWE-corr)** | **T** | **x,y,z {mm}** | **Anatomical Label** |
| ***p*** | | | | | |
| 0 | 705 | 0.001 | 5.38 | 9 45 -26 | Right superior frontal gyrus, orbital part |
| 0.005 | 131 | 0.005 | 5.02 | -2 51 23 | Left superior frontal gyrus, medial part |
| 0 | 468 | 0.005 | 5.01 | 0 -14 11 | Left thalamus |
| 0.003 | 165 | 0.01 | 4.85 | -8 29 -23 | Left gyrus rectus |
| 0.031 | 12 | 0.014 | 4.78 | 54 15 29 | Right inferior frontal gyrus, opercular part |
| 0.02 | 32 | 0.014 | 4.77 | -41 14 -17 | Left inferior frontal gyrus |
| 0.002 | 195 | 0.016 | 4.75 | 5 -98 -3 | Right cuneus |
| 0.035 | 8 | 0.021 | 4.68 | 20 -15 -11 | Right sub-lobar |
| 0.018 | 39 | 0.023 | 4.65 | -53 -11 42 | Left postcentral gyrus |
| 0.036 | 7 | 0.028 | 4.61 | 59 -11 29 | Right postcentral gyrus |
| 0.04 | 4 | 0.044 | 4.49 | -26 48 18 | Left middle frontal gyrus |
| ***Thought*** | | | | | |
| 0 | 1616 | 0 | 5.6 | 8 44 -26 | Right gyrus rectus |
| 0 | 519 | 0 | 5.51 | -2 51 23 | Left superior frontal gyrus, medial |
| 0.013 | 57 | 0.002 | 5.2 | 54 15 29 | Right inferior frontal gyrus, opercular part |
| 0 | 558 | 0.003 | 5.12 | 0 -14 11 | Left thalamus |
| 0.003 | 167 | 0.003 | 5.09 | -53 -11 41 | Left postcentral gyrus |
| 0.009 | 88 | 0.006 | 4.98 | -41 14 -17 | Left inferior frontal gyrus |
| 0.001 | 318 | 0.006 | 4.96 | -8 -84 -17 | Left lingual gyrus |
| 0.019 | 36 | 0.012 | 4.82 | 59 -11 29 | Right postcentral gyrus |
| 0.013 | 57 | 0.013 | 4.79 | -26 48 20 | Left superior frontal gyrus |
| 0.032 | 11 | 0.018 | 4.71 | 20 -17 -11 | Right brainstem |
| 0.033 | 10 | 0.027 | 4.62 | 39 15 -20 | Right inferior frontal gyrus |
| 0.037 | 6 | 0.032 | 4.57 | 47 5 -6 | Right superior temporal gyrus/insula |
| 0.033 | 10 | 0.033 | 4.56 | 26 3 -17 | Right subcallosal gyrus/amygdala |
| 0.039 | 5 | 0.038 | 4.53 | 44 8 -14 | Right superior temporal gyrus |
| 0.043 | 2 | 0.04 | 4.52 | 35 18 -30 | Right superior temporal pole |
| ***Internalizing*** | | | | | |
| 0.015 | 50 | 0.005 | 4.99 | -6 -83 -17 | Left lingual gyrus |
| 0.028 | 16 | 0.008 | 4.91 | 54 15 29 | Right inferior frontal gyrus, opercular part |
| 0.036 | 7 | 0.031 | 4.58 | -2 51 23 | Left superior frontal gyrus, medial |
| 0.027 | 19 | 0.033 | 4.57 | 9 44 -26 | Right superior frontal gyrus, orbital part |
| ***Externalizing*** | | | | | |
| 0 | 742 | 0 | 6.37 | -44 32 17 | Left inferior frontal gyrus, triangular part |
| 0 | 11849 | 0 | 6.26 | 0 54 18 | Left superior frontal gyrus, medial |
| 0 | 954 | 0 | 5.85 | -41 8 -17 | Left superior temporal gyrus/insula |
| 0 | 393 | 0 | 5.58 | -51 -12 36 | Left postcentral gyrus |
| ***Cluster*** | | ***Peak*** | | | |
| **p(FWE-corr)** | **k** | **p(FWE-corr)** | **T** | **x,y,z {mm}** | **Anatomical Label** |
| ***Externalizing*** | | | | | |
| 0 | 1110 | 0.001 | 5.39 | 5 3 45 | Right supplementary motor area |
| 0 | 566 | 0.001 | 5.32 | 42 9 -17 | Right superior temporal gyrus |
| 0.002 | 198 | 0.003 | 5.16 | -45 8 30 | Left precentral gyrus/inferior frontal gyrus |
| 0.005 | 131 | 0.004 | 5.08 | 27 35 41 | Right middle frontal gyrus |
| 0.006 | 111 | 0.004 | 5.04 | 18 -32 -3 | Right thalamus |
| 0.014 | 55 | 0.005 | 5.02 | 20 -17 -11 | Right brainstem |
| 0.004 | 146 | 0.006 | 4.97 | 26 -42 -12 | Right fusiform gyrus/parahippocampal gyrus |
| 0.001 | 307 | 0.007 | 4.94 | 2 -11 9 | Right thalamus |
| 0.022 | 28 | 0.009 | 4.88 | 56 -24 48 | Right postcentral gyrus |
| 0.015 | 51 | 0.009 | 4.87 | 53 14 29 | Right inferior frontal gyrus, opercular part |
| 0.006 | 118 | 0.01 | 4.86 | 50 -9 45 | Right precentral gyrus |
| 0.014 | 55 | 0.016 | 4.74 | -33 -83 -18 | Left fusiform gyrus |
| 0.016 | 45 | 0.017 | 4.73 | 56 -11 29 | Right precentral gyrus |
| 0.021 | 30 | 0.02 | 4.69 | 15 60 -6 | Right superior frontal gyrus, orbital part |
| 0.04 | 4 | 0.027 | 4.62 | 42 24 12 | Right inferior frontal gyrus, triangular part |
| 0.027 | 19 | 0.028 | 4.61 | -12 -30 -5 | Right brainstem |
| 0.031 | 13 | 0.03 | 4.59 | 57 -23 -9 | Right middle temporal gyrus |
| 0.03 | 14 | 0.033 | 4.56 | -60 -20 -9 | Left middle temporal gyrus |
| 0.037 | 6 | 0.037 | 4.54 | -18 -14 -11 | Left extra-nuclear |
| 0.043 | 2 | 0.041 | 4.51 | -47 -32 -6 | Left temporal lobe |
| 0.037 | 6 | 0.042 | 4.51 | 6 -92 -6 | Right lingual gyrus |
| 0.046 | 1 | 0.047 | 4.48 | 6 -20 59 | Right supplementary motor area |
| 0.043 | 2 | 0.047 | 4.48 | 35 -39 44 | Right supramarginal gyrus |
| 0.046 | 1 | 0.048 | 4.47 | 42 17 32 | Right inferior frontal gyrus, opercular part |
| 0.046 | 1 | 0.049 | 4.46 | 12 62 14 | Right superior frontal gyrus, medial |
